# Host variation in type I interferon signaling genes (*MX1*), *CCR5*Δ*32*, and MHC class I alleles in treated HIV+ non-controllers predict viral reservoir size

**DOI:** 10.1101/2021.10.31.466670

**Authors:** David A. Siegel, Cassandra Thanh, Eunice Wan, Rebecca Hoh, Kristen Hobbs, Tony Pan, Erica A. Gibson, Deanna L. Kroetz, Peter W. Hunt, Jeffrey Martin, Frederick Hecht, Christopher Pilcher, Jeffrey Milush, Maureen Martin, Mary Carrington, Satish Pillai, Michael P. Busch, Mars Stone, Claire N. Levy, Meei-Li Huang, Pavitra Roychoudhury, Florian Hladik, Keith R. Jerome, Hans-Peter Kiem, Timothy J. Henrich, Steven G. Deeks, Sulggi Lee

**Affiliations:** Department of Medicine, Division of HIV, Infectious Diseases & Global Medicine, University of California San Francisco, 995 Potrero Avenue, San Francisco, CA 94110, USA; Institute of Human Genetics, University of California San Francisco, CA 94143, USA; Department of Bioengineering and Therapeutic Sciences, University of California San Francisco, CA 94158; Department of Medicine, Division of Experimental Medicine, University of California San Francisco, CA 94110, USA; Department of Biostatistics & Epidemiology, University of California San Francisco, CA 94158, USA; Basic Science Program, Frederick National Laboratory for Cancer Research, National Cancer Institute, Frederick, MD and Laboratory of Integrative Cancer Immunology, Center for Cancer Research, National Cancer Institute, Bethesda, MD USA; Ragon Institute of MGH, MIT and Harvard, Cambridge, MA, USA; Vitalant Blood Bank, San Francisco, CA 94118 USA; Department of Obstetrics and Gynecology, University of Washington, WA, 98105 USA; Department of Laboratory Medicine and Pathology, University of Washington, Seattle WA 98195, USA; Vaccine and Infectious Disease Division, Fred Hutchinson Cancer Research Center, Seattle WA 98109, USA

**Keywords:** HIV reservoir, host genetics, type I interferon, MHC class I, CCR5

## Abstract

**Objective:** Prior genomewide association studies have identified variation in MHC Class I alleles and *CCR5*Δ*32* as genetic predictors of viral control, especially in “elite” controllers, individuals who remain virally suppressed in the absence of therapy.

**Design:** Cross-sectional genomewide association study.

**Methods:** We analyzed custom whole exome sequencing and direct HLA typing from 202 ART-suppressed HIV+ non-controllers in relation to four measures of the peripheral CD4+ T cell reservoir: HIV intact DNA, total (t)DNA, unspliced (us)RNA, and RNA/DNA. Linear mixed models were adjusted for potential covariates including age, sex, nadir CD4+ T cell count, pre-ART HIV RNA, timing of ART initiation, and duration of ART suppression.

**Results:** Previously reported “protective” host genetic mutations related to viral setpoint (e.g., among elite controllers) were found to predict smaller HIV reservoir size. The HLA “protective” B*57:01 was associated with significantly lower HIV usRNA (q=3.3×10^−3^), and among the largest subgroup, European ancestry individuals, the *CCR5*Δ*32* deletion was associated with smaller HIV tDNA (p=4.3×10^−3^) and usRNA (p=8.7×10^−3^). In addition, genomewide analysis identified several SNPs in *MX1* (an interferon stimulated gene) that were significantly associated with HIV tDNA (q=0.02), and the direction of these associations paralleled *MX1* gene eQTL expression.

**Conclusions:** We observed a significant association between previously reported “protective” MHC class I alleles and *CCR5*Δ*32* with the HIV reservoir size in non-controllers. We also found a novel association between *MX1* and HIV total DNA (in addition to other interferon signaling relevant genes, *PPP1CB*, *DDX3X*). These findings warrant further investigation in future validation studies.

## Introduction

Although antiretroviral therapy (ART) prolongs life, it does not fully restore health, as evidenced by persistently high levels of immune activation ^[1]^ and increased rates of non-AIDS-related mortality ^[2]^ observed in HIV-infected compared to uninfected individuals ^[3–6]^. Persistent HIV may contribute to ongoing inflammation, immune activation, and subsequent clinical outcomes, even during effective ART ^[5–8]^. Identifying host genetic predictors of the HIV reservoir in ART-suppressed individuals may shed light on novel (and potentially modifiable) targets to reduce the HIV reservoir and inflammation- and immune activation-associated adverse effects on long-term morbidity and mortality.

Most prior host genetic HIV studies have focused on identifying variants associated with viral setpoint, e.g., among “elite controllers”, HIV+ individuals able to maintain viral suppression in the absence of therapy^[9–18]^. These studies identified several key single nucleotide polymorphisms (SNPs) in the human Major Histocompatibility Complex (MHC), or human leukocyte antigen (HLA)-B and -C regions as well as deletions in the C-C chemokine receptor type 5 gene (*CCR5*Δ*32*)^[19–22]^ and a SNP in the HLA complex 5 (*HCP5*) gene^[10]^. However, whether *residual* viral control during *treated* HIV disease – i.e., “the HIV reservoir” – is influenced by the same genetic variants is unknown. We performed custom whole exome sequencing among HIV non-controllers in relation to four measures of the peripheral CD4+ T cell HIV reservoir: cell-associated “intact” DNA^[23]^, total DNA, unspliced RNA, and RNA/DNA (**Figure S1**). We found that previously reported “protective” HLA-B*57:01^[10, 17]^ and *CCR5*Δ*32*^[20, 21, 24]^ mutation were associated with smaller HIV reservoir size. Genomewide analyses demonstrated several novel associations with SNPs in interferon signaling-associated genes (*MX1*, *PPP1CB*, *DDX3X*) and total HIV DNA reservoir size. Gene set enrichment analysis identified several interferon signaling-associated genes to significantly predict intact HIV DNA levels in the largest subgroup, Europeans.

## Methods

### Study participants

HIV+ non-controllers who initiated ART during chronic (>2 years) or early (<6 months) HIV infection were sampled from the UCSF SCOPE and Options cohorts (**Table S1**). Inclusion criteria were laboratory-confirmed HIV-1 infection, availability of 10×10^6^ cryopreserved PBMCs, and plasma HIV RNA levels below the limit of assay quantification (<40 copies/mL) for at least 24 months at the time of biospecimen collection. HIV “controllers,” individuals with a history of undetectable viral load in the absence of therapy for at least 1 year prior to the specimen collection date^[25–27]^, were excluded. The estimated date of detected infection (EDDI) was calculated for each study participant to determine recency of infection in relation to ART start date using detailed clinical HIV diagnostic test results, using the Infection Dating Tool (https://tools.incidence-estimation.org/idt/)^[28]^. Additional exclusion criteria were potential factors that might influence HIV reservoir quantification, including recent hospitalization, infection requiring antibiotics, vaccination, or exposure to immunomodulatory drugs in the six months prior to sampling timepoint. The research was approved by the UCSF Committee on Human Research (CHR), and all participants provided written informed consent.

### Custom whole exome host DNA sequencing

Genomic DNA was extracted (AllPrep Universal Kit, Qiagen, Hilden, Germany) from negatively selected CD4+ T cells from cryopreserved PBMCs (StemCell, Vancouver, Canada). Targeted exome capture was performed with custom addition of 50 Mb regulatory regions (Roche NimbleGen, Wilmington, MA), sequencing libraries were generated and then run on the Illumina HiSeq 2000 system (Illumina, San Diego, CA). The custom regions included 50 kb upstream and 50 kb downstream of 442 candidate genes related to cell cycle regulation, HIV host restriction factors, and HIV-host integration, which were selected based on Gene Ontology (GO) Consortium experimental evidence codes (EXP, IDA, IPI, IMP, IGI, IEP) (**Table S2**).

### HLA typing

Direct HLA typing was performed from extracted genomic DNA following the PCR-SSOP (sequence-specific oligonucleotide probing) typing and PCR-SBT (sequence based typing) protocols recommended by the 13th International Histocompatibility Workshop^[29, 30]^. Locus-specific primers were used to amplify a total of 25 polymorphic exons of HLA-A & B (exons 1-4), C (exons 1-5), E (exon 3), DPA1 (exon 2), DPB1 (exons 2-4), DQA1 (exon 1-3), DQB1 (exons 2-3), DRB1 (exons 2-3), and DRB3, 4, 5 (exon 2) genes with Fluidigm Access Array (Fluidigm, Singapore) and sequenced on an Illumina MiSeq sequencer (Illumina, San Diego, USA). HLA alleles and genotypes are called using the Omixon HLA Explore (version 2.0.0) software (Omixon, Budapest, Hungary).

### HIV reservoir quantification from peripheral CD4+ T cells

The HIV reservoir largely consists of “defective” virus that harbors mutations prohibiting the production of infectious virus^[31, 32]^. There is currently no “gold standard” for measuring the HIV reservoir. Therefore, we estimated the frequency of HIV “intact” DNA using a ddPCR-based assay to quantify the size of the potentially “replication-competent” reservoir^[23, 33, 34]^. We also measured HIV total DNA (quantifies both defective and intact HIV) and unspliced RNA (quantifies full-length HIV RNA) using an HIV-1 LTR-specific quantitative polymerase chain reaction (qPCR) TaqMan assay^[35]^. DNA and RNA were simultaneously dual extracted using the AllPrep Universal Kit (Qiagen, Hilden, Germany). HIV tDNA and usRNA were then quantified in triplicate reaction wells using a 7-point standard curve (1–10,000 copies/second). To estimate the frequency of “intact” HIV DNA, five regions on the HIV genome were interrogated in a multiplex ddPCR assay^[23]^. Droplet generation and thermocycling were performed according to manufacturer instructions. To determine potentially replication-competent (“intact”) HIV genomes, the number of positive droplets for 3 targets per assay were quantified. Two targets in a housekeeping gene (*RPP30)* were used to quantify all cells, and a target in the T cell receptor D gene (*TRD)* was used to identify all non-T cells, to normalize the HIV copy numbers/10^6^ CD4+ T cells. A DNA shearing index (DSI) (using *RPP30*) was then used to calculate the estimated number of intact HIV genomes after correcting for shearing.

### Data processing and quality control

The bcbio bioinformatics pipeline^[36]^ was used to perform DNA alignment, which included the Burroughs-Welcome Aligner (BWA) tool^[37]^ and the GenomeAnalysisToolkit (GATK) HaplotypeCaller joint variant calling method^[38]^. Reads were initially mapped to reference genome b37, then transposed to human genome assembly GRCh38 using Picard tools^[39]^. SNPs and insertions or deletions (indels) were then filtered by variant quality score recalibration (VQSR) using GATK^[40]^. The whole genome data analysis toolset, PLINK^[41]^, was then used to validate the chromosomal sex of each individual, filter out individuals with excessive heterozygosity, and SNPS violating Hardy-Weinberg equilibrium (HWE) at a p-value threshold of 1×10^−8^. The VCFtools suite of functions were then used to summarize data, run calculations, convert data, and filter out data, and convert data, and filter out relevant SNPs^[42]^.

The GENESIS analysis pipeline^[43]^ was used to analyze the relatedness and ancestries of the individuals in the study. All individuals were determined to be unrelated (kinship estimates <0.05) aside from one pair of siblings, so one sibling was randomly removed from the study. The remaining 199 unrelated individuals had diverse and mixed ancestries (**Figure 1**). We accounted for population stratification in the total population by (1) including a genetic effects term with a genetic relatedness matrix (GRM), (2) by including the first five PCs as covariates in the multivariate models, and by (3) performing sensitivity analyses among the largest subgroup, Europeans.

**Figure 1.**
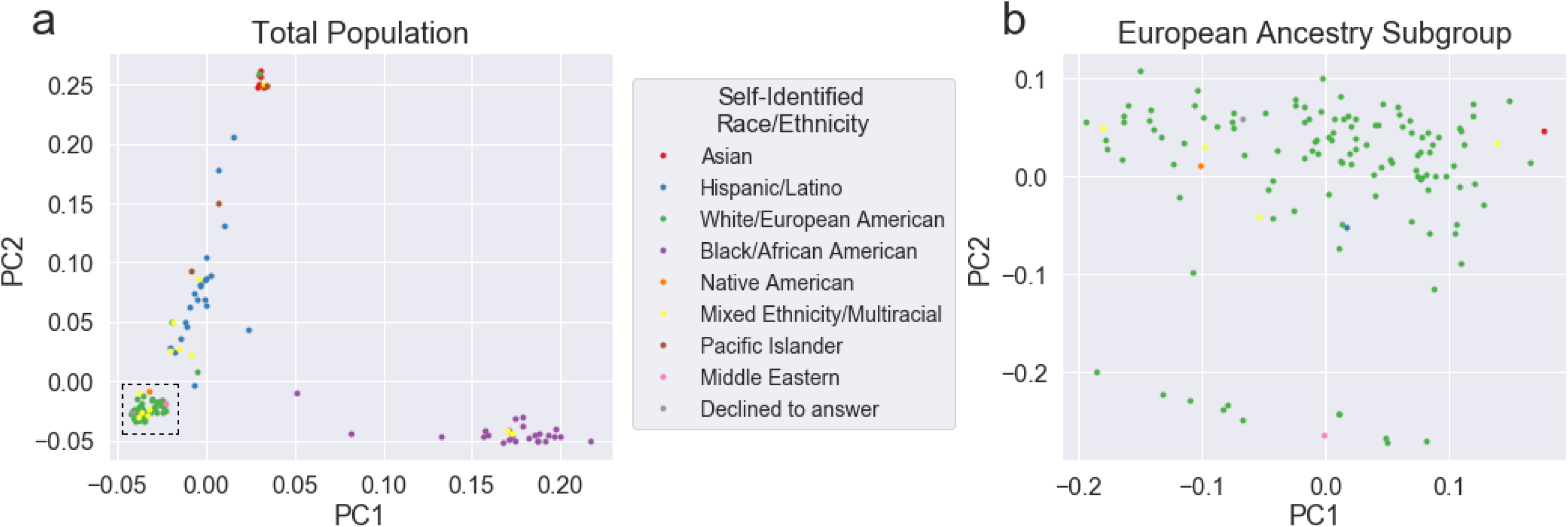
Genetic principal component analysis (PCA) plots of the full study population (with legend with self-identified race/ethnicity) (a) and of the largest homogenous ancestral population, males of mostly European ancestry (b). Recalculated European ancestry male PCA plot is shown in panel b, from lower left dashed box in panel a.

### Single SNP common variant analyses

Individual SNP associations were calculated with GENESIS “assocTestSingle”. For HIV total DNA, unspliced RNA, RNA/DNA, and intact DNA, respectively, the outcome variables were: log_10_((DNA copies/10^6^ CD4+ T cells + offset)); log_10_((RNA copies/10^6^ CD4+ T cells + offset)); log_10_((RNA copies/106 CD4+ T cells + offset) / (DNA copies/106 CD4+ T cells + offset)); log_10_((Intact DNA copies/10^6^ CD4+ T cells + offset)). The offsets for RNA and DNA counts were given by the smallest nonzero measured values of RNA and DNA, respectively, to avoid divergences in the logarithm. Final covariates in multivariate models were sex, timing of ART initiation (**Figure S2**), nadir CD4+ T cell count (**Figure S3**), and the first 5 PCs. Pre-ART viral load (**Figure S4**) and duration of ART suppression (**Figure S5**) were not associated with HIV reservoir size nor improved the fit of the final models. A Gaussian link function was used, and a GRM was included with results filtered for SNPs with MAF ≥5%. SNP annotations were then obtained using Annovar^[44]^.

### Gene-based rare variant analyses

Gene level multi-SNP associations were calculated with the GENESIS software package “assocTestAggregate” function implementing the variant Set Mixed Model Association Test (SMMAT)^[45]^ for alleles with MAF<5% with weights following the beta distribution parameters of *a*_1_=1 and *a*_2_=25^[46]^. The same covariates, GRM, and regression family were used as for the individual SNP associations. Outcomes were quantile-normalized to follow a normal distribution. Gene regions were defined according to UCSC hg38 assembly^[47]^.

Gene set enrichment analyses (GSEA) were performed using the Molecular Signatures Database (MSigDB)^[48, 49]^. For all gene set analyses, introns and flanking regions of *±*50kb were included in the SMMAT p-value calculations for each gene to account for potential regulatory regions and SNPs with smaller effects. GSEAPreranked was run with default parameters on the SMMAT gene-level *−log*10(*P*).

### HLA analysis

Multivariate regression models were fit using the python statsmodels OLS function^[50]^ with covariates for sex, timing of ART initiation, nadir CD4+ T cell count, and 3 genetic PCs.

## Results

### Study population

A total of 202 HIV-infected ART-suppressed individuals from the UCSF SCOPE and Options HIV+ cohorts were included in the study. Consistent with our San Francisco-based HIV patient population, participants were mostly male (94%) with median age of 46 (**Table S1**). Participants had a median of 5.1 years of ART suppression, a median nadir CD4+ T cell count=341 cells/mm^3^, and pre-ART HIV RNA=5.1 log_10_ copies/mL. The majority of study participants reported White/European American ethnicity (63%), and the remainder reported Black/African American (12%), Hispanic/Latino (11%), Mixed Ethnicity/Multiracial (6%), Asian (4%), Pacific Islander (1.5%), Native American (<1%), and Middle Eastern (<1%) ethnicities. Most study participants (N=147) had highly detailed clinical test results to be able to calculate their estimated date of detected infection (EDDI), but a subset of 55 study participants only had self-reported data regarding date of ART initiation in relation to date of HIV seroconversion. For these individuals (all of whom initiated ART prior to widespread guidelines for initiating ART at the time of HIV diagnosis^[51]^), we mean-imputed values assuming ART initiation starting after 2 years from infection. This estimation is supported by prior data from our cohort and others demonstrating that the HIV reservoir size is relatively stable after 2 years of infection^[52–56]^. Overall results for all final models were unchanged when performing sensitivity analyses excluding those with imputed values for timing of ART initiation.

### Earlier ART initiation and lower nadir CD4+ T cell count were associated with smaller HIV reservoir, and HIV reservoir measures were correlated with each other

Consistent with prior work^[32, 57, 58]^, earlier ART initiation was associated with significantly smaller HIV reservoirs (tDNA, usRNA, intact DNA) (**Figure S2**), while lower nadir CD4+ T cell count was associated with larger HIV reservoir (tDNA, usRNA, intact DNA, RNA/DNA) (**Figure S3**). Pre-ART viral load (**Figure S4**) and duration of ART suppression (**Figure S5**) were not associated with HIV reservoir size. Although usRNA was correlated with both tDNA intact DNA (**Figure S6a-b**), tDNA was not associated with intact DNA (**Figure S6c**).

### HLA “protective” B*57:01 and “risk” C*07 alleles were associated with smaller and larger HIV reservoir sizes, respectively

Using a Benjamini-Hochberg false discovery rate (FDR) adjusted q<0.05 threshold^[59]^, we examined previously reported protective (B*57:01, B*27:05, B*14, C*08:02, B*52, and A*25) and risk (B*35 and C*07) alleles for viral setpoint in untreated HIV+ controllers^[17]^ and found a “protective” association with HLA-B*57:01 and usRNA (β=−1.5, q=3.3×10^−3^), with a similar trend observed with tDNA (β=−1.6, q=0.13). Similarly, previously reported HLA-C*07 “risk” allele also demonstrated a “risk” trend (larger reservoir size) in our European subgroup (tDNA: β=0.76, q=0.072, and usRNA: β=0.41, q=0.10). Further analyses employing a composite HLA variable did not identify statistically significant associations (**Tables S3-S6**).

### *CCR5*Δ*32* was associated with smaller HIV reservoir size

Deletions in the C-C chemokine receptor type 5 gene (*CCR5*Δ*32*) have previously been shown to be associated with HIV viral control in the absence of therapy^[20, 21, 24]^. Among individuals of European ancestry (where *CCR5*Δ*32* is more commonly observed), the presence *CCR5*Δ*32* was associated with smaller HIV reservoir size (tDNA: β=−1.3, p=4.3×10^−3^; usRNA: β=−0.78, p=8.7×10^−3^), with a similar trend observed in the total population (tDNA: β=−0.86, p=0.045; usRNA: β=−0.41, p=0.12), In addition, the previously reported long noncoding RNA variant which regulates differential CCR5 expression (rs1015164)^[22]^, was found to be significantly associated with smaller HIV reservoir size in Europeans (usRNA: β=−0.39, p=0.027), which reached near-statistical significance in the total population as well (usRNA: β=− 0.30, p=0.051),

### Genomewide association analysis identified several SNPs in *MX1* associated with larger and smaller HIV reservoir sizes, paralleling predicted *MX1* gene expression

A total of 1,279,156 variants from 23,733 genes were included in the final analysis from 199 study participants whose sequencing data passed quality control metrics (**Figure S7**). Final models demonstrated lambda genomic inflation factor^[60]^ values close to 1, supporting adequate adjustment for possible bias due to population stratification (ancestry) (**Figure 2**).

**Figure 2.**
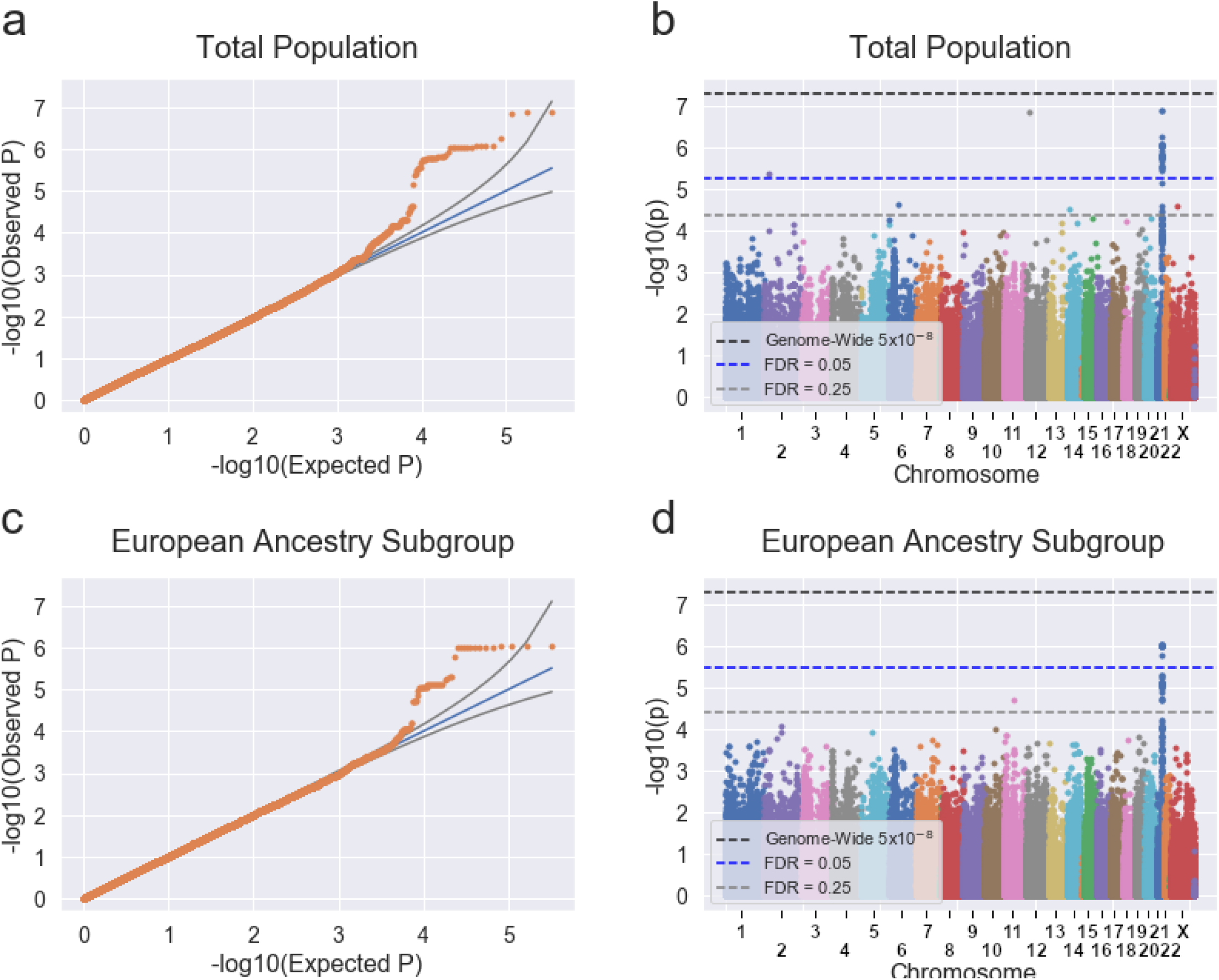
Quantile-quantile (QQ) plots (a, c) and Manhattan plots (b, d) of the total study population (a-b) and of European ancestry (c-d). QQ plots: the blue line represents the expected −log10 p-values while the black lines denote the expected error bars. Manhattan plots: the horizontal black line delineates a traditional conservative genome-wide significance of p-value of 5×10^−8^, while less conservative Benjamini-Hochberg false discovery rate (FDR) statistical significance of q=0.05 is shown as the horizontal blue line (q=0.25 is shown in grey).

The strongest genomewide associations were observed with HIV total DNA reservoir measures (**Tables 1 and S7**). In particular, 44 SNPs in linkage disequilbrium (LD) in the human interferon-inducible myxovirus resistance 1 gene, *MX1*, also known as *MXA*[61, 62] were significantly associated with tDNA (all q<0.03). *MX1* is closely related to *MX2* (*MXB*), which encodes a well-known potent host restriction factor that inhibits HIV-1 infection^[63–65]^. We then compared the directionality of the SNP hits with previously reported whole blood eQTL data at these loci^[66–68]^ and found the *MX1* SNPs associated with larger total HIV DNA reservoir sizes seemed to be in eQTL regions predicting increased *MX1* expression and vice versa (**Table S7**). We also observed two additional SNPs significantly associated with HIV tDNA, the first in *PPP1CB* (encodes Protein Phosphatase 1 Catalytic Subunit Beta, which reduces antiviral potency of MX2 against HIV-1^[65]^, q=0.03) and the second in *LRMP* (encodes Lymphoid-Restricted Membrane Protein, which plays a critical role in the delivery of peptides to MHC class I molecules^[69]^, q=0.03) (**Table 1**). Additional SNPs that showed non-statistically significant trends with HIV tDNA were in *DDX3X* (DEAD-box helicase 3 X-linked, regulates the production of type I interferons^[70]^, q=0.17) and *AKAP6* (A-Kinase Anchoring Protein 6, binds to protein kinase A regulatory subunits, a critical signaling pathway associated with HIV latency reversal and T cell proliferation^[71, 72]^, q=0.20). Among Europeans, *OSBP* (oxysterol-binding protein, associated with HIV-1 infection of monocyte-derived macrophages from highly-exposed seronegative individuals^[73]^, q=0.14), showed a non-significant trend with HIV tDNA.

**Table 1.**
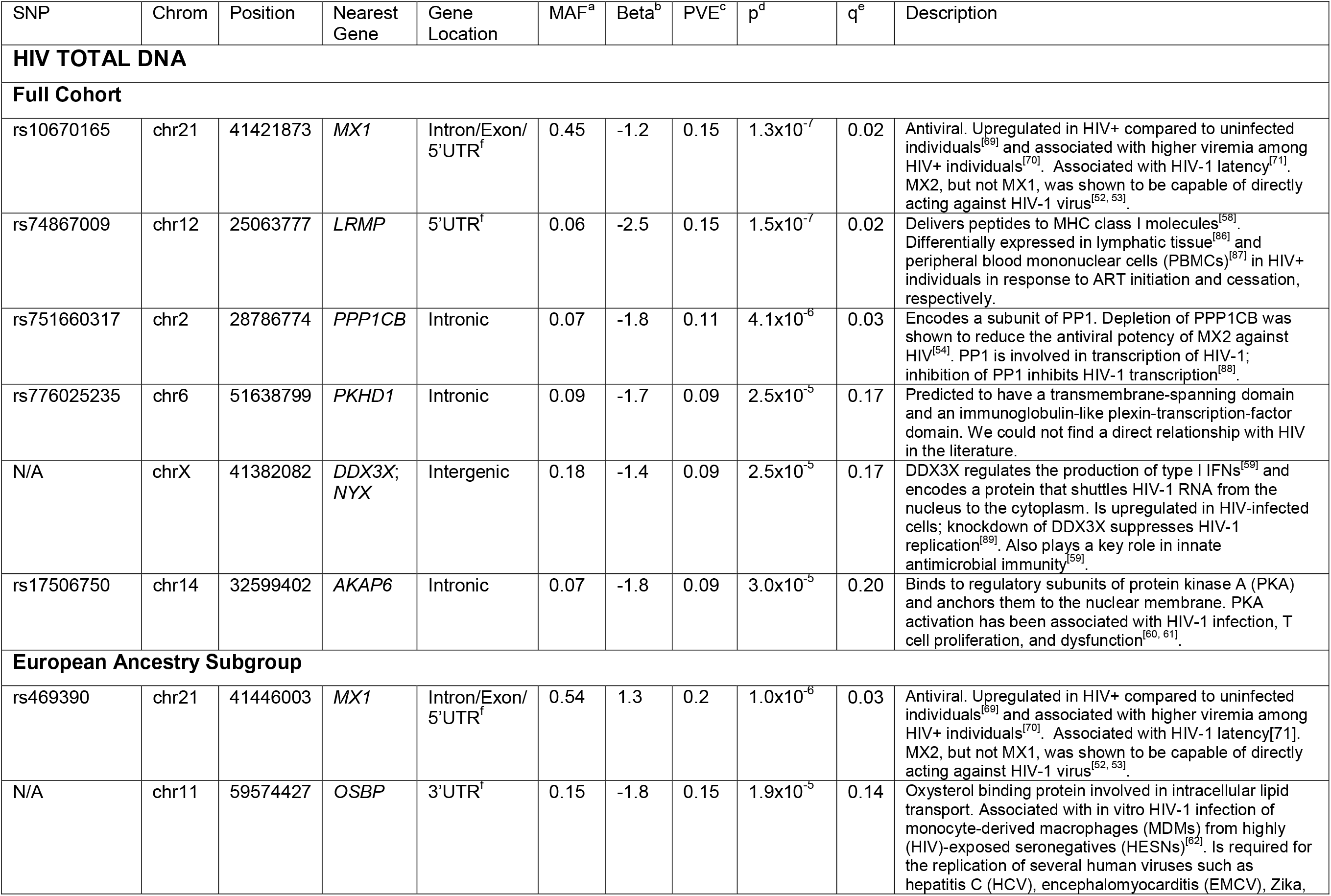
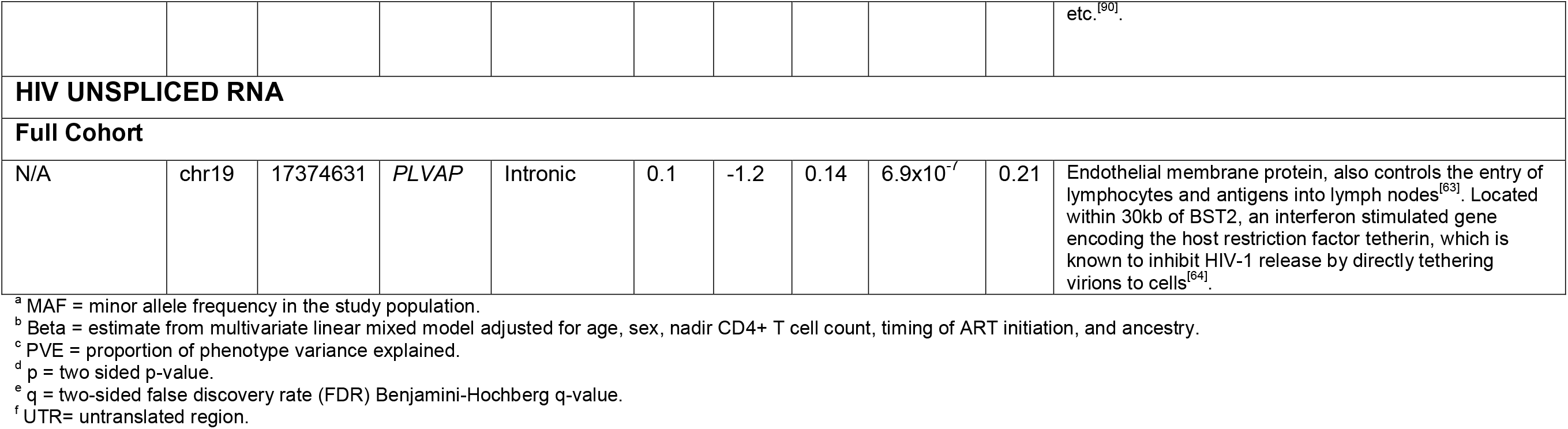
Single nucleotide polymorphisms (SNPs) associated with HIV total DNA (upper table) and unspliced RNA (lower table) in the total study ation. Two additional SNPs were significantly associated with total HIV DNA in the subpopulation of European ancestry (middle table). For in which there were several SNPs in linkage disequilibrium (LD), the top SNP for each gene is shown, with the full list of SNPs shown in S7.

Although not statistically significant, a SNP in *PLAVP* (protein regulating lymphocyte migration into lymph nodes^[74]^, q=0.21), lying <30 kilobases upstream of *BST2* (tetherin, an HIV host restriction factor^[75]^) demonstrated a non-significant trend with usRNA (**Table 1**). No SNPs met statistical significance in association with HIV intact DNA or RNA/DNA ratio.

### Gene set enrichment analysis demonstrated several interferon signaling-associated genes associated with intact HIV DNA

We then performed multi-SNP analyses to identify genes associated with HIV reservoir size. GSEA identified several interferon signaling-associated genes (e.g., *IFITM1*, *IFITM3*, *APCS*, *IFITM2*, *FCN3*, *FCN1*, *GSN*, *TRIM59*, *SNX3*, *TRIM25*, *PTX3*, *TRIM11*, *TRIM8*, *MID2*, *TRIM5*, *IFNA2*) in the gene set called “Negative Regulation of Viral Entry into Host Cell,” to significantly predict HIV intact DNA (q=0.03) (**Figure 3, Table S8**). Several other gene sets showed non-significant trends with HIV reservoir size (**Figure 3, Table S8**), including gene sets related to interferon-induced STAT signaling and intact DNA, glycosylation and tDNA, and retroviral transcription and usRNA.

**Figure 3.**
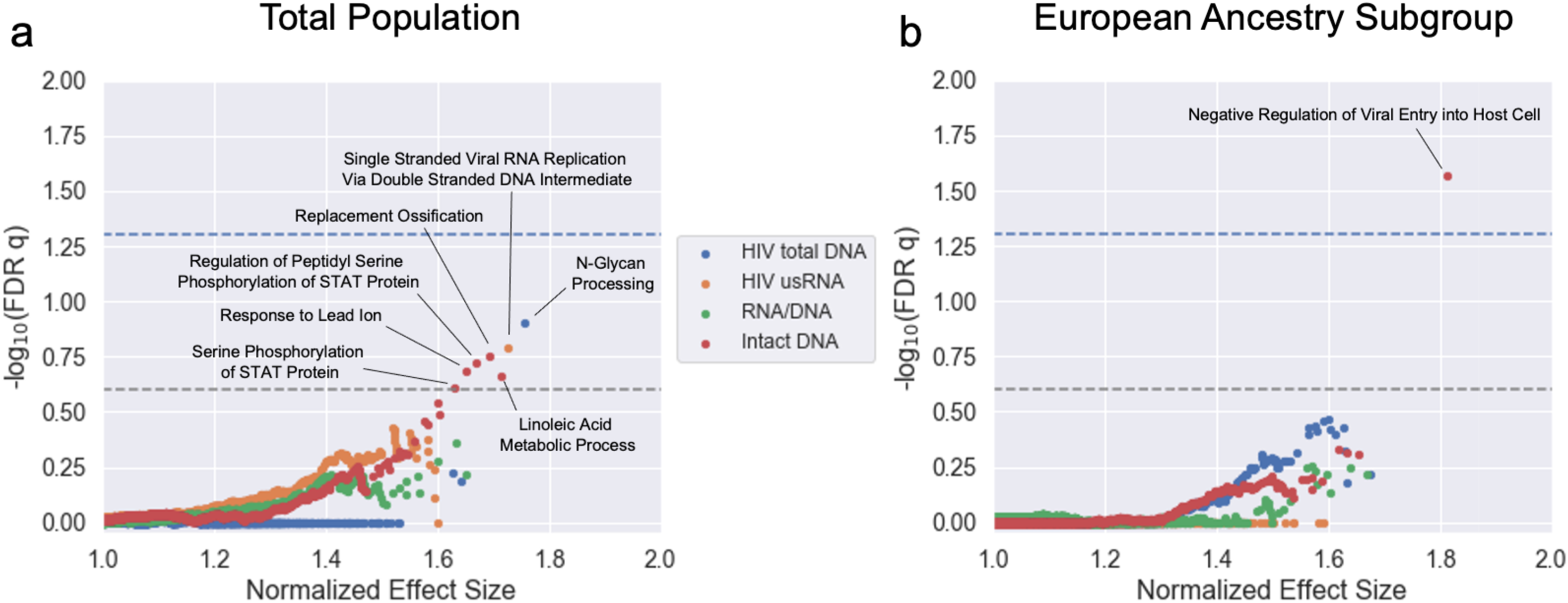
Gene set enrichment analysis (GSEA) was used to identify associations between genes, ordered by p-value under a host multi-SNP rare variant (minor allele frequency, MAF, <5%) analysis using HIV total DNA, unspliced RNA, RNA/DNA, and intact DNA as outcomes; and biological processes gene sets, in the total study population (a) and among European ancestry (b). Horizontal dashed lines represent the GSEA Benjamini-Hochberg false discovery rate (FDR) statistical significance level of q=0.05 (blue) and q=0.025 (grey), respectively.

## Discussion

HIV eradication remains a critical goal in reducing long-term morbidity and mortality among all PLWH since life-long viral suppression does not appear to fully restore health, as evidenced by persistently high levels of immune activation^[2]^ and high rates of mortality^[8]^ in HIV-infected compared to healthy individuals. ART is also expensive, carries long-term risk of toxicity, and poses major challenges in being able to be accessible to a global population for decades^[76]^. HIV cure clinical trials to date have yielded disappointing results^[77–82]^. Novel approaches are needed to better target potential immunologic pathways that help maintain the HIV reservoir.

Our study is the first host genomic study to evaluate several measures of the peripheral HIV reservoir in HIV+ non-controllers, including quantification of HIV intact DNA, an estimate of the putative “replication-competent” reservoir by droplet digital PCR^[23, 33, 83]^. We also performed direct HLA typing of 25 polymorphic exons of HLA-A & B, C, E, DPA1, DPB1, DQA1, DQB1, DRB1, and DRB3,4,5. We performed individual SNP and gene-based analyses including detailed clinical covariate data such as timing of ART initiation, one of the strongest clinical predictors of HIV reservoir size, which enhanced the fit of our final multivariate models, potentially allowing us to detect otherwise difficult-to-discern genetic effects. Unlike prior genomic studies that have primarily focused on the ~1% of the HIV+ population able to suppress virus in the absence of therapy (“elite controllers”), we focused on HIV+ ART-suppressed non-controllers, which make up the majority of people living with HIV (PLWH). We found that prior significant HLA and *CCR5*Δ*32* genetic associations predicting viral setpoint among HIV+ elite controllers^[10, 19–22]^ in our study, predicted the HIV reservoir size. We also identified several additional (uninvestigated) host genetic variants associated with the HIV reservoir (signals that might have been attenuated in a study population enriched for “stronger” genetic effects, such as HLA and/or *CCR5*Δ*32*).

The most striking finding from our SNP-based analysis was the identification of several SNPs in *MX1*, which encodes for a potent antiviral factor which inhibit replication of several RNA viruses, including influenza A and measles, and DNA viruses, such as hepatitis B^[84]^. *MX1* expression has also been shown to be upregulated in HIV+ vs. HIV-uninfected individuals^[85]^, in HIV+ individuals with high vs. low viremia^[86]^, and with HIV-1 latency in latently-infected cell lines^[87]^. Genomewide (i.e., DNA-based) results cannot directly infer directionality of gene function without further functional studies. However, for our top hit SNPs in *MX1*, we compared the directionality of the SNP hits with previously reported whole blood eQTL data at these loci^[66–68]^ and found that the *MX1* SNPs associated with larger total HIV DNA reservoir sizes seemed to be in eQTL regions predicting increased *MX1* expression and vice versa. However, determining whether a single variant is responsible for both genomewide and eQTL signals in a locus can be challenging. Nonetheless, as a further query, we performed colocalization analysis, an *in silico* method to integrate GWAS and eQTL results to calculate a *probability* of whether a SNP is causal for both an eQTL and disease trait^[88]^, but only found a 1% probability that the *MX1* top SNPs are causally linked to both gene expression and HIV reservoir size. Gene-based analyses also identified several interferon signaling-associated genes (within the “negative regulation of viral entry into host cell” gene set) that significantly predict intact HIV DNA (**Table S8**), but as these genes were not in eQTL regions, the putative directionality of these associations could not be further queried. Additional functional genomic validation, e.g., CRISPR-Cas9 editing of primary human T cells^[89]^, is needed to further investigate the potential role of *MX1* (and other interferon signaling genes) in HIV persistence.

We also found that the previously reported “protective” HLA-B*57:01 and *CCR5*Δ*32* mutation were associated with smaller HIV reservoir size in our study. These findings suggest that immune pathways that control viral setpoint during untreated disease may also play a role in the maintenance of the HIV reservoir during treated infection. It is also possible that identified genetic variants may have variable “penetrance”^[90]^ – e.g., the genetic variants that may drive “elite control” might similarly, but less obviously, influence HIV persistence in treated non-controllers.

There are limitations to our study that deserve mention. Although the HIV reservoir has been shown to be relatively stable over time^[58, 91, 92]^, our cross-sectional design provides a “snapshot” of the HIV reservoir after a median of 5.1 years of ART suppression and may not reflect genetic associations with reservoir decay. Second, as is characteristic of many U.S.-based HIV+ cohorts, our San Francisco-based population consisted mostly of males of European ancestry. Population stratification is a critically important potential bias in any multiethnic genomic study. Thus, we approached this in least three ways using well-established methods to adjust for population stratification bias^[43, 93]^: first by calculating principal components and including these as covariates in the final models, second by including a genetic relatedness matrix (GRM) in the models, and finally by performing stratified analyses, focusing on the largest homogenous subpopulation (individuals with European ancestry). Overall, the findings observed in the European ancestral group did not overlap with the non-European (e.g., African-American) subgroup (**Table S9**). Thus, it is critical that these results be replicated in larger studies, especially those including women and individuals from different ethnic backgrounds. Third, the majority of the HIV reservoir persists in lymphoid tissues, not in the periphery^[94]^. Although recent data suggests that the tissue compartment largely reflects (and is the likely source of) the peripheral compartment^[52, 95, 96]^, it will be important to determine whether the results from our study are generalizable to the tissue HIV reservoir. Fourth, intact HIV DNA represents the putative replication-competent reservoir. Although we observed several genes that were significantly associated with intact HIV DNA in the gene set enrichment analyses, individual genes did not meet statistical significance. This may be due to the challenge in estimate the frequency of intact and/or replication-competent HIV when in fact, the majority of the HIV reservoir consists of defective HIV. For these reasons, quantitative outgrowth assays and assays to measure intact HIV DNA often result in many low/ zero values compared to total HIV DNA, which has a larger dynamic range^[83, 97, 98]^. In our study, HIV intact DNA was undetectable in over half of our measured samples, while for example, total DNA was measurable in 95% of samples (**Figure S6**). With so many samples below the limit of detection for intact DNA, the statistical power to detect SNP associations is much lower for this assay than for the other HIV reservoir assays included in our study. However, when we were able to enhance the ability to detect an association by performing the gene set enrichment analyses (essentially, a method that aggregates several rare variants into immunologically relevant “gene sets” to test for an association with HIV reservoir size), we observed several statistically significant associations with HIV intact DNA in the total population (STAT signaling, critical for regulating the innate and adaptive immune responses) and among individuals of European ancestry, the largest subgroup with the greatest statistical power (“negative regulation of viral entry into host cell” – which included several interferon signaling genes) (**Figure 3, Table S8**).

Our findings are in contrast to two recent genomewide studies of the HIV reservoir, which did not identify an association with *MX1*, HLA-B*57:01, or *CCR5*Δ*32*, nor reported similar findings to each other^[99, 100]^. The first study performed GWAS microarray genotyping from 797 HIV+ treated individuals (194 with whole exome sequencing data), included several longitudinal measures of HIV total DNA from peripheral blood mononuclear cells (PBMCs), and imputed HLA alleles (from genotypes), but did not observe any significant associations with HLA alleles, *CCR5*Δ*32*, or SNPs^[99]^. The second study included 207 HIV+ treated individuals and performed GWAS microarray (no HLA or *CCR5*Δ*32* typing) and included measures of HIV tDNA and usRNA from peripheral CD4+ T cells. They reported a significant association between tDNA and a SNP in *PTDSS2* (phosphatidylserine synthase 2) at genomewide p<5×10^−8^, which was not statistically significant in our analysis.^[100]^ These differences highlight a particular challenge in performing genomic studies, let alone for studies of the HIV reservoir size (which can be measured in several different ways as total HIV DNA, HIV unspliced RNA, HIV intact DNA, etc.) from different biospecimens (PBMCs, CD4+ T cells). Add to this the use of different study designs (untreated HIV+ controllers, treated HIV+ non-controllers – and those treated during chronic versus acute infection), and analytic methods (multivariate models with or without key clinical covariates such as timing of ART initiation, nadir CD4+ T cell count, etc.) and a lack of understanding regarding the exact mechanism by which the genetic code is expressed from DNA to RNA to protein (which also varies by cell type and within different tissues^[101]^) – then differences between these three small studies might fall within the expected range of variability. Given the polygenic nature of the host immune response, the contribution of host genetics in predicting HIV reservoir size might vary widely, leading to variable results when comparing small genomic studies. Prior genomewide association studies of HIV progression during untreated disease explain ~13% of the variability in viral load, with strong genetic predictors such as HLA and *CCR5*Δ*32*^[102]^. Using a tool for genome-wide complex trait analysis (GCTA)^[103]^, we calculated the heritability of our total HIV DNA phenotype to be up to 0.78, but the error bars were large (+/−0.94). This suggests that unknown host genetic loci might play a significant role in determining the size of the HIV reservoir – but that there is a high degree of variability in that estimate. For these reasons, we are careful to describe our study results within the limits of a genomewide association study identifying potential novel DNA variants related to HIV persistence (e.g., only highlighting *MX1* as it lies with an eQTL region but the others for which there are no functional data, we do not) and again emphasize the need for functional studies to pursue the novel hypotheses identified from our discovery-based study. Our findings may vary from the two prior published studies due to several differences in (1) study design (cross-sectional, only including HIV+ ART-suppressed non-controllers), (2) statistical modeling (detailed clinical covariates for timing of ART initiation, nadir CD4+ T cell count, etc.), (3) HIV reservoir quantification (e.g., HIV total DNA, unspliced RNA, and intact DNA), (4) sampling (CD4+ T cells, not bulk PBMCs), (5) host genomic assays (custom whole exome sequencing instead of GWAS microarray, and direct HLA typing instead of imputed HLA alleles), and (6) approach to handling potential population stratification bias (at least three methods - principal component analysis, genetic related matrix methods, and sensitivity analyses restricted to ancestral subgroups).

Using carefully selected ART-suppressed HIV non-controllers, we performed custom whole exome sequencing and direct HLA typing, quantified several measures of the peripheral HIV reservoir, and fit multivariate models adjusted for clinical and demographic covariates that influenced the size of the HIV reservoir, and found that the previously reported “protective” HLA-B*57:01 and the favorable *CCR5*Δ*32* (during untreated disease) were associated with smaller HIV reservoir size. Genomewide analyses identified several SNPs in *MX1*, a type I interferon stimulated gene, were significantly associated with total HIV DNA, which correlated with predicted *MX1* eQTL gene expression, and HIV intact DNA were associated with several interferon signaling-associated genes in gene set enrichment analyses. Our findings support a surprising role of the innate immune response (e.g., genes involved in interferon signaling^[104–107]^) in maintaining the HIV reservoir during long-term suppressive ART. These findings support recent studies demonstrating that measures of cell-associated HIV RNA correlate with time to viral rebound^[106, 108–111]^. Perhaps host genes driving immediate antiviral responses play a major role in maintaining the HIV reservoir if the “transcriptionally active” reservoir is indeed a major source of the “rebound-competent” reservoir.” Additional studies are needed to functionally validate these findings, especially among more diverse patient populations, including female and non-European HIV+ patients.

## Acknowledgements

The authors wish to acknowledge the participation of all the study participants who contributed to this work as well as the clinical research staff of the SCOPE and Options who made this research possible. All funders had no role in study design, data collection and analysis, decision to publish, or preparation of the manuscript. All authors provided critical feedback in finalizing the report. SAL conceived and designed the study with critical feedback from SGD, PWH, TH, DLK, KRJ, and SP. SGD, JM, FH, CP, RH, and SAL coordinated the collection, management, and quality control processes for the cohort clinical data and SGD, JM, FH, CP, MPB, MS provided biospecimens. SAL and EW performed the whole exome sequencing assays. SAL, CT, JM, KB, TP, EAG performed participant sample processing, SAL and EW performed the whole exome sequencing, and DAS and SAL performed quality control analyses and the genomic association analyses for the study. SAL, CT, and KH performed the qPCR HIV reservoir assays (total DNA, unspliced RNA) in the lab of TH. CNL and MLH performed the ddPCR HIV reservoir assay (intact DNA) in the labs of FH, KRJ, and HPK. PR, DAS, TJH, and SAL analyzed these HIV reservoir data in relation to host genomic and clinical phenotype data. MM and MC performed the HLA typing to determine HLA alleles for the analyses. DAS and SAL wrote the report. All authors provided critical feedback in finalizing the manuscript.

