## Supplemental Tables and Figures for "Host variation in type I interferon signaling genes (*MX1*), *CCR5*Δ*32*, and MHC class I alleles in treated HIV+ non-controllers predict viral reservoir size"

**Figure S1.** Study participant sample selection flowchart. Specific inclusion and exclusion criteria are listed for each selection step and for HIV reservoir measure analysis.

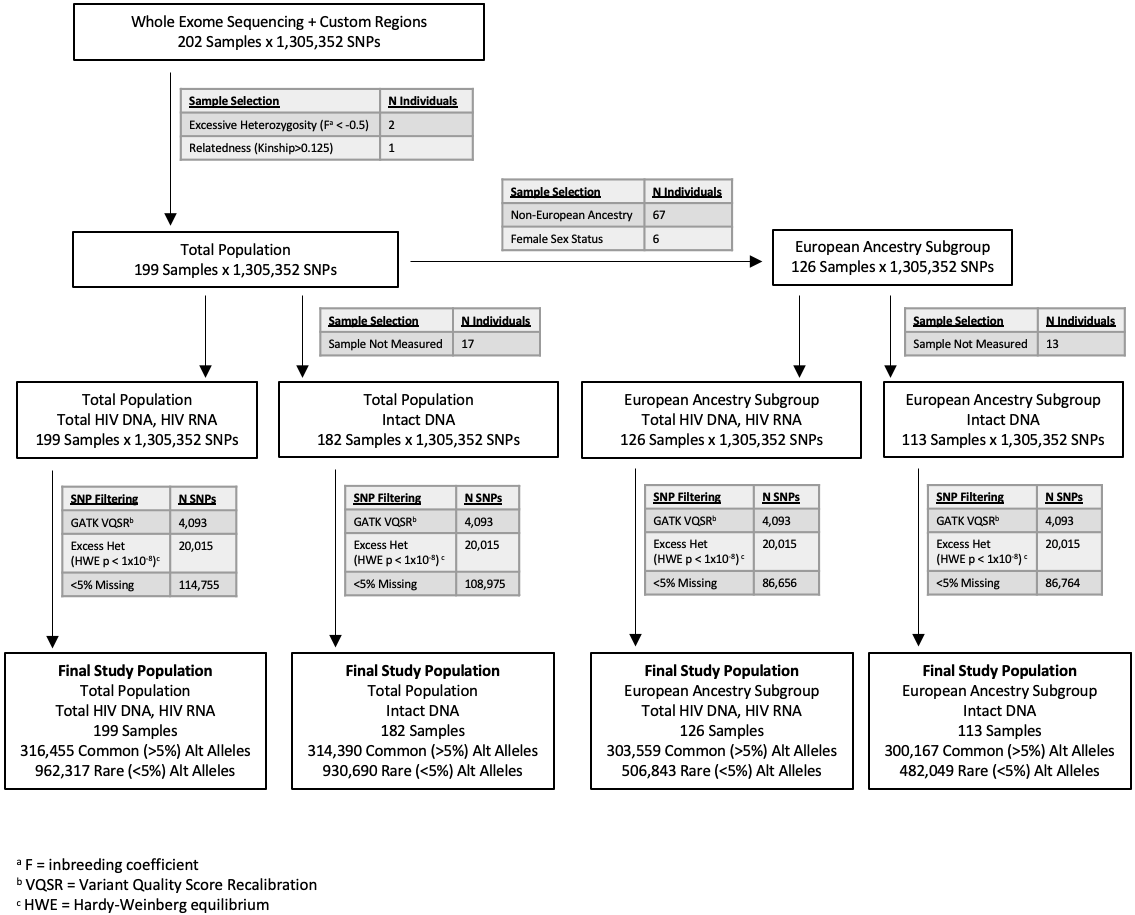

**Table S1.** Descriptive statistics for the study population of 202 HIV-infected ART-suppressed non-controllers. Median frequencies (with interquartile ranges) are shown below unless otherwise specified.

| **Descriptive Characteristic** | **Total**  (N=202) | **Early-Treated^a^**  (N=57) | **Later-Treated^a^**  (N=145) |
| --- | --- | --- | --- |
| Male (%)^b^ | 192 (95%) | 57 (100%) | 135 (93%) |
| Age (years) | 46 (13) | 43 (12) | 47 (13) |
| Nadir CD4+ T cell count (cells/mm3) | 341 (259) (N=201) | 537 (333) | 300 (201) (N=144) |
| Maximum pre-ART HIV RNA (log_10_copies/mL)^c^ | 5.1 (0.9) (N=197) | 5.6 (0.7) (N=54) | 5.0 (0.9) (N=143) |
| Duration of ART suppression (years)^c^ | 5.1 (4.1) (N=200) | 6.0 (4.3) (N=55) | 4.7 (4.0) |
| Timing of ART initiation (years)^c^ | 2.0 (4.5) (N=190) | 0.23 (0.19) | 3.4 (4.3) (N=133) |
| HIV intact DNA (log_10_copies/10^6^ CD4+ T cells) | 1.2 (1.1) (N=182) | 1.9 (1.0) (N=46) | 1.2 (0.7) (N=136) |
| HIV total DNA (log_10_copies/10^6^ CD4+ T cells) | 1.0 (1.3) | 0.4 (1.3) | 1.2 (1.2) |
| HIV unspliced RNA (log_10_copies/10^6^ CD4+ T cells) | 3.2 (0.8) | 3.0 (0.8) | 3.3 (0.7) |
| HIV RNA/DNA | 2.3 (1.0) | 2.4 (1.0) | 2.2 (0.9) |

^a^ Early-treated = Individuals who initiated ART within 6 months of the date of detected HIV infection; later-treated = Individuals who initiated ART after 6 months of date of detected HIV infection.

^b^ Absolute frequencies (with percent).

^c^ Adjudicated data from clinical records and patient history were available for a total of: 201 (nadir CD4+ T cell count), 197 (pre-ART HIV RNA), 200 (duration of ART suppression until date of specimen), and 190 (timing from estimate date of HIV infection to ART initiation) of the total 202 study participants.

**Table S2.** List of custom genes included in the whole exome sequencing. For each, custom sequencing included 50 kb upstream and 50 kb downstream of the gene start and end loci.

|  | List of Genes |
| --- | --- |
| Immune Checkpoint Inhibitor Genes | *PDCD1, CD274, PDCD1LG2, PTPN6, PTPN11, ZAP70, CD247, LCK, CSK, AKT1, AKT2, AKT3, PIK3R1, PIK3R2, PIK3R3, PIK3R4, PIK3R5, PIK3R6, FOXO3, MTOR, CCR5, HLA-A, HLA-B, HLA-C, KIR2DL1, KIR2DL2, KIR2DL3, KIR2DL4, KIR2DL5A, KIR2DL5B, KIR3DL1, KIR3DL2, KIR3DL3, KIR2DS1, KIR2DS2, KIR2DS3, KIR2DS4, KIR2DS5, KIR3DS1* |
| Host Restriction Factor Genes | *APOBEC3A, APOBEC3B, APOBEC3C, APOBEC3D, APOBEC3F, APOBEC3G, APOBEC3H, BST2, CD74, CDKN1A, CH25H, CNP, CTR9, EIF2AK2, HERC5, IFITM1, IFITM2, IFITM3, IRF1, IRF7, ISG15, LGALS3BP, MOV10, MX2, PAF1, RNASEL, RSAD2, RTF1, SAMHD1, SERINC3, SERINC5, SLFN11, SUN2, TNFRSF10A, TRIM5, TRIM11, TRIM14, PML, TRIM21, TRIM22, TRIM26, TRIM28, TRIM32, BRD4* |
| HIV Integration Genes | *BACH2, MKL2, STAT5B, CYTH1, RPTOR, TAOK1, TNRC6B, VPS45, PRKCB, MKL1, MAP4, PAK2, NSD1, KIAA0319L, SLC30A7, SHOC2, NUMA1, CDKN1B, ARID2, NFATC3, SMARCE1, EPS15L1, GATAD2A, HORMAD2, ALG12, BTBD9, OSBPL3, UBE2H, FNBP1* |
| GO Cell Cycle Genes | *ZNF503, AIF1, IFNG, IL12B, IL12A, PID1, TAL1, CDC27, CDK6, CDK4, BTG3, CDK14, CDK9, SLC9A3R1, CCPG1, RAD17, RGCC, LILRB1, LEF1, CENPF, TCF7L2, CDC7, FZR1, GATA3, CCNO, CENPE, CDKN2A, MCPH1, MAP2K1, GNB2L1, ESX1, GAS2L1, PES1, CIB1, NKX3-1, MADD, MX2, ING4, TIMELESS, THBS1, TP53, ADARB1, CDKN1B, XRCC2, PCBP4, DCLRE1B, ZNF703, YY1AP1, SOX2, TGFB2, ZW10, CDKN2D, CTBP1, ATM, DTL, MAD2L1, ZAK, TARDBP, MLF1, CENPC, MECOM, INTS3, RAB6C, ZWILCH, IRF1, CDKN3, PRR11, MASTL, CXCL8, INHA, NOTCH2, NABP1, KIF14, MAD2L1BP, KIAA0101, PNPT1, UHMK1, GADD45GIP1, ZBTB49, STK11, CENPW, SMC1A, PUM1, WDR6, KIF18B, RASSF1, PRDM5, KLLN, TGFB1, HRAS, MYC, NABP2, OVOL2, NBN, PHF13, UBE2L3, BIRC2, PRCC, CEP250, BAP1, CHFR, SMAD3, CENPT, FER, APC, SON, BOP1, GADD45A, USP3, FOXO4, DDIT3, GRK5, CDKN2C, CDKN2B, PTPRC, INHBA, NEK4, BCL2L1, ZWINT, WNT9A, IRF6, CALR, TIPIN, PML, TBRG1, CDKN1A, SIRT2, BEX2, KNTC1, USP17L2, RBM38, NEK11, BIN1, RB1, MEN1, HPGD, TCF3, ETS1, LRP6, GMNN, ANKRD17, BMP4, BMP2, ZNF143, TAF6, SOX9, DAPK3, RPS15A, TRIM32, RNF167, RARA, THAP5, C6orf89, RPS6KA2, PTPRK, FOXM1, RPRD1B, CYLD, THAP1, ASUN, PIM3, TXLNG, ZNF268, SLC25A33, GBF1, BRINP1, SIRT1, BMP7, TRIM21, TBX3, AFAP1L2, TP73, ALOX15B, SFRP1, PLK1, HMGA2, MCIDAS, E4F1, NLRP2P, BRCA1, KDM8, DUSP3, EIF4E, ASNS, ABCB1, LATS2, MIF, CCNK, USP2, USP22, CCNB1, TRIM35, TERF1, FBXL15, PLK3, RRP8, INSM1, FOXA1, MDM2, PLK2, XPC, TFAP4, ARPP19, NES, UHRF2, TPD52L1, DDB1, PKN2, POLE, PIM2, MED25, ACVR1, RCC1, HMGN5, MLXIPL, SETMAR, SCRIB, MYBBP1A, SPHK1, DLG1, FOXC1, DGKZ, PKD2, EIF4EBP1, FOXE3, USP37, CDCA5, NUDT16, NPPC, RPS6KB1, CDC25C, CDC25B, ERCC2, PTPN3, ERCC3, SPDYA, PRKCA, CCND1, ZNF16, USH1C, SOX11, GATA6, HINFP, PHF8, CACUL1, LATS1, FGF10, BIRC5, CHEK2, CCNY, TAF2, DAB2IP, CRLF3, BRSK2, ACVR1B, TFDP3, GAS1, DBF4B, PTEN, KCNH5, CDC73, ID2, PHB2, DDX3X, LCMT1, SUSD2, PPP1R1C, INO80, VPS4B, H2AFY, GEN1, TPR, RAC1, XRCC3, RAD51C, TP63, E2F1, RAB11A, DACT1, C10orf99, UBE2E2, USP47, ZC3H12D, SLFN11, RCC2, LSM11, STXBP4, NEK10, STOX1, BRD7, RAD51B, FHL1, BRD4, LSM10, ATP2B4, DYNC1LI1, JADE1, PCID2, PLK5, WIZ, C8orf4, PTPN6, ZNF655, KANK2, ANAPC15, MUC1, CRADD, PPP2R5C, SOX4, CASP2, PIDD1, MDM4, CYP27B1, FBXO31, FOXN3, RPA2, TAOK3, RPS27L, CDK5RAP3, RFWD3, BLM, ATF2, PRKDC, DUX4, RHNO1, RINT1, BRCC3, RBBP8, TAOK2, BABAM1, MAPKAPK2, BRE, UIMC1, BRSK1, WAC, CDC14B, FAM175A, TAOK1, CHEK1, NEK6, CLSPN* |

Definitions of the above custom gene set lists are included below:

Immune Checkpoint Inhibitor Genes. Previously reported immune checkpoint inhibitor genes associated with PD1-signaling in Ref (45).

Host Restriction Factor Genes. Previously reported host restriction factor genes in Ref. (46)

HIV Integration Genes. Previously reported genes into which HIV integrates 3 or more times among the 5 patients in Ref (47).

Gene Ontology (GO, Biologic Processes) Cell Cycle Genes. Genes with "cell cycle" annotation either directly or inferred, or inferred annotation of "virus", "virion", or "viral", in MSigDB (Ref (48)).

**Figure S2.** HIV total DNA, unspliced RNA, RNA/DNA, and intact DNA were associated with timing of ART initiation. Spearman correlation and p-values are shown in the inset boxes.

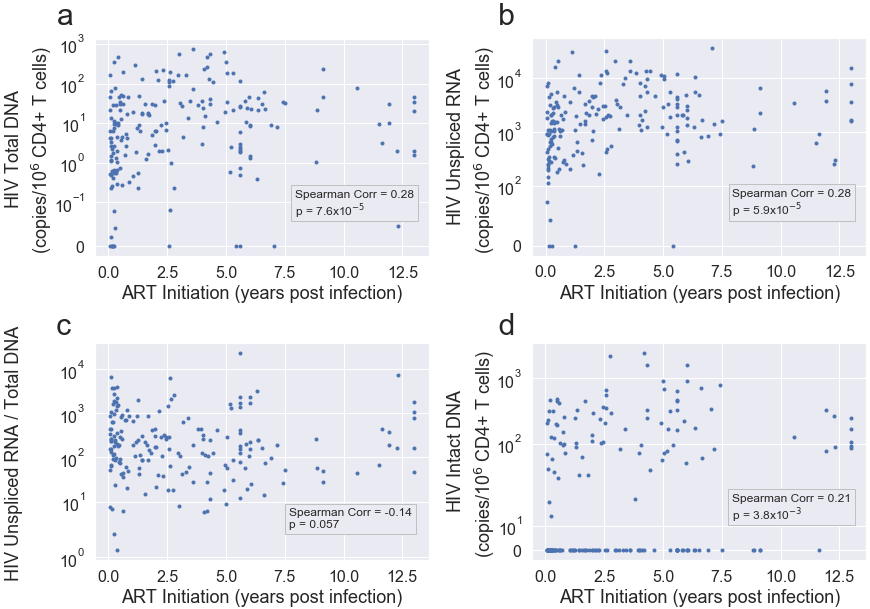

**Figure S3.** Nadir CD4+ T cell counts were associated with measures of the HIV reservoir: total DNA (a), unspliced RNA (b), RNA/DNA (c), and intact DNA (d). Spearman correlation and corresponding p-value are shown in the inset boxes.

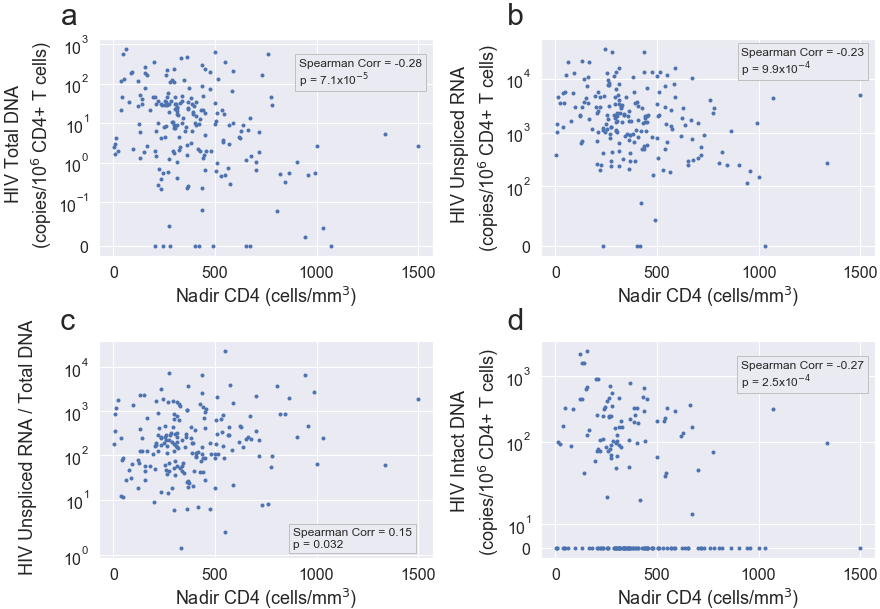

**Figure S4.** Maximum pre-ART viral loads prior ART initiation (and at least 1 year prior to specimen collection for these analyses) were not associated with measures of the HIV reservoir: total DNA (a), unspliced RNA (b), RNA/DNA (c), and intact DNA (d). Spearman correlation and corresponding p-value are shown in the inset boxes.

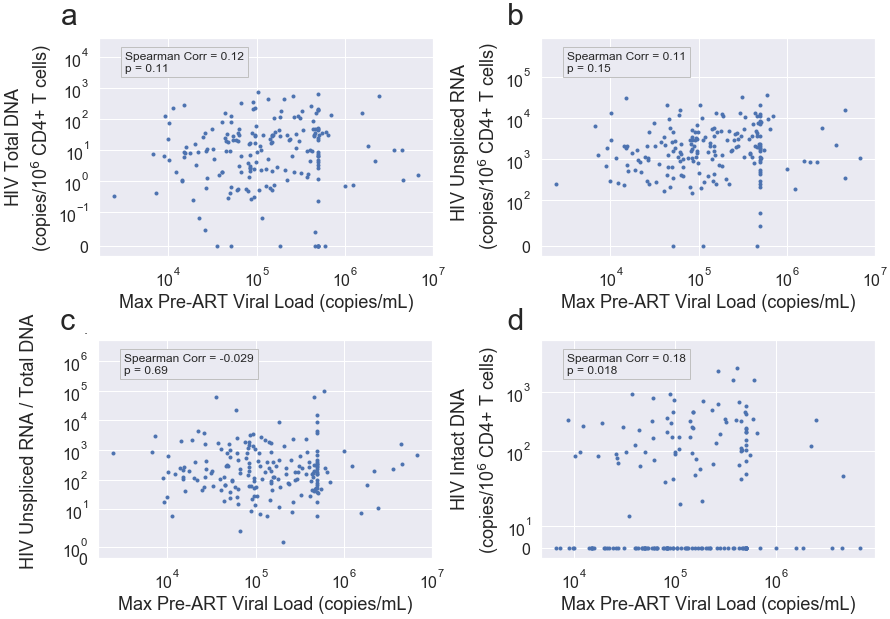

**Figure S5.** Duration of ART suppression at the time of specimen collection was not associated with measures of the HIV reservoir: total DNA (a), unspliced RNA (b), RNA/DNA (c), and intact DNA (d). Spearman correlation and corresponding p-value are shown in the inset boxes.

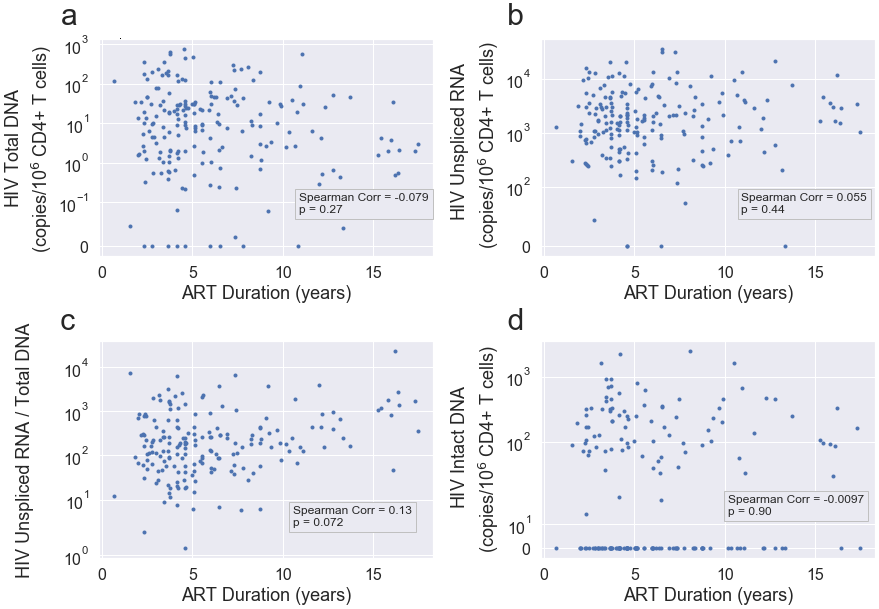

**Figure S6.** Measures of HIV total DNA (a) and HIV intact DNA (b) were statistically significantly correlated with HIV unspliced RNA, but HIV total DNA was not correlated with intact DNA (c). Spearman correlation and p-values are shown in the inset boxes.

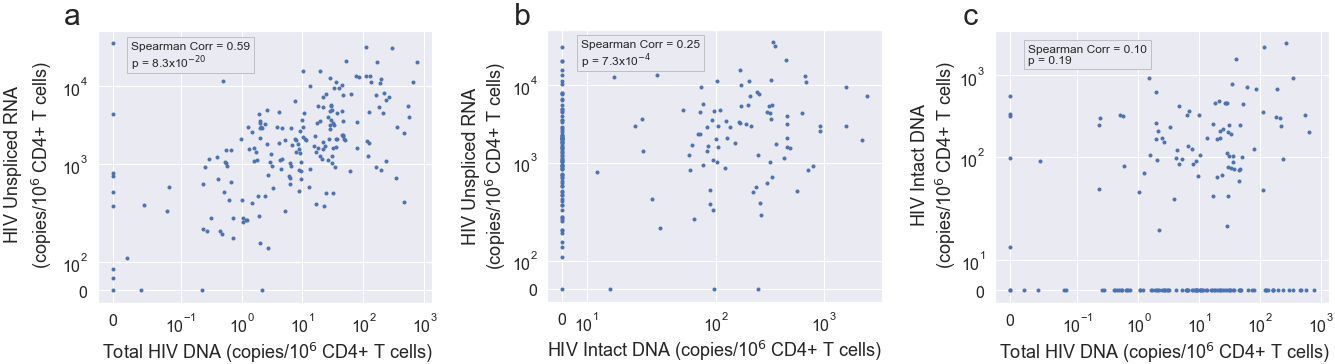

**Figure S7.** Distribution of the total number of host variants (a), as well as all exonic variants (b) sequenced in the study population.

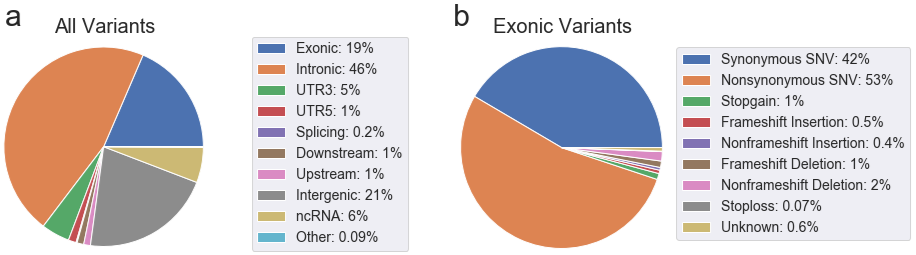

**Table S3.** HLA allele associations with HIV total DNA (copies/10^6^ CD4+ T cells).

| **HIV Total DNA** | | | | | | | | | |
| --- | --- | --- | --- | --- | --- | --- | --- | --- | --- |
|  |  | **Full Cohort** | | | | **European Subgroup** | | | |
|  | HLA Allele^a^ | MAC^b^ | Beta^c^ | p^d^ | q^e^ | MAC^b^ | Beta^c^ | p^d^ | q^e^ |
| 1 | B*57:01 | 10 | -1.6 | *0.032 | 0.13 | 9 | -1.44 | *0.041 | 0.12 |
| 2 | B*27:05 | 11 | 0.79 | 0.26 | 0.36 | 4 | 0.37 | 0.73 | -- |
| 3 | B*14 | 24 | 0.41 | 0.36 | 0.41 | 19 | 0.083 | 0.86 | 0.86 |
| 4 | C*08:02 | 23 | 0.23 | 0.62 | 0.62 | 17 | -0.15 | 0.77 | 0.86 |
| 5 | B*52 | 3 | 1.7 | 0.2 | -- | 2 | 1.5 | 0.32 | -- |
| 6 | A*25 | 7 | 0.96 | 0.27 | 0.36 | 5 | 1.4 | 0.13 | -- |
| 7 | B*35 | 46 | -0.81 | *0.021 | 0.13 | 32 | -0.6 | 0.11 | 0.22 |
| 8 | C*07 | 107 | 0.4 | 0.14 | 0.28 | 74 | 0.76 | *0.012 | 0.072 |
| 9 | Composite Variable | N/A | 0.32 | 0.066 | 0.18 | N/A | 0.13 | 0.48 | 0.72 |

^a^ Alleles 1-6 were previously designated as “protective” alleles, while alleles 7-8 were defined as “risk” alleles, in the International HIV Controllers genomewide association study (16). The “Composite Variable” combines each of the above alleles, counting each protective allele as +1 and each risk allele as -1.

^b^ We list minor allele count (MAC), rather than MAF, due to the multiallelic nature of these variants. Alleles with MAC<=5 are greyed out to highlight low observed frequencies in our study population.

^c^ Beta estimate from multivariate linear mixed model adjusted for age, sex, nadir CD4+ T cell count, timing of ART initiation, and ancestry.

^d^ Two-sided p-values; p<0.05 are annotated with asterisks.

^e^ Two-sided false discovery rate (FDR)-adjusted Benjamini-Hochberg q-values.

**Table S4.** HLA allele associations with HIV unspliced RNA (copies/10^6^ CD4+ T cells).

| **HIV Unspliced RNA** | | | | | | | | | |
| --- | --- | --- | --- | --- | --- | --- | --- | --- | --- |
|  |  | **Full Cohort** | | | | **European Subgroup** | | | |
|  | HLA Allele^a^ | MAC^b^ | Beta^c^ | p^d^ | q^e^ | MAC^b^ | Beta^c^ | p^d^ | q^e^ |
| 1 | B*57:01 | 10 | -1.5 | *4.2x10^-4^ | 0.0033 | 9 | -1.5 | *5.3x10^-4^ | 0.0032 |
| 2 | B*27:05 | 11 | 0.39 | 0.35 | 0.7 | 4 | 0.13 | 0.85 | -- |
| 3 | B*14 | 24 | 0.041 | 0.88 | 0.88 | 19 | -0.047 | 0.88 | 0.88 |
| 4 | C*08:02 | 23 | -0.078 | 0.77 | 0.88 | 17 | -0.14 | 0.67 | 0.8 |
| 5 | B*52 | 3 | 0.56 | 0.47 | -- | 2 | -0.21 | 0.83 | -- |
| 6 | A*25 | 7 | 0.15 | 0.77 | 0.88 | 5 | 0.43 | 0.48 | -- |
| 7 | B*35 | 46 | -0.44 | *0.032 | 0.13 | 32 | -0.24 | 0.31 | 0.62 |
| 8 | C*07 | 107 | 0.2 | 0.21 | 0.56 | 74 | 0.41 | *0.034 | 0.1 |
| 9 | Composite Variable | N/A | 0.058 | 0.57 | 0.88 | N/A | -0.056 | 0.64 | 0.8 |

^a^ Alleles 1-6 were previously designated as “protective” alleles, while alleles 7-8 were defined as “risk” alleles, in the International HIV Controllers genomewide association study (16). The “Composite Variable” combines each of the above alleles, counting each protective allele as +1 and each risk allele as -1.

^b^ We list minor allele count (MAC), rather than MAF, due to the multiallelic nature of these variants. Alleles with MAC<=5 are greyed out to highlight low observed frequencies in our study population.

^c^ Beta estimate from multivariate linear mixed model adjusted for age, sex, nadir CD4+ T cell count, timing of ART initiation, and ancestry.

^d^ Two-sided p-values; p<0.05 are annotated with asterisks.

^e^ Two-sided false discovery rate (FDR)-adjusted Benjamini-Hochberg q-values.

**Table S5.** HLA associations with HIV unspliced RNA/ total DNA.

| **HIV RNA/DNA** | | | | | | | | | |
| --- | --- | --- | --- | --- | --- | --- | --- | --- | --- |
|  |  | **Full Cohort** | | | | **European Subgroup** | | | |
|  | HLA Allele^a^ | MAC^b^ | Beta^c^ | p^d^ | q^e^ | MAC^b^ | Beta^c^ | p^d^ | q^e^ |
| 1 | B*57:01 | 10 | 0.067 | 0.91 | 0.91 | 9 | -0.091 | 0.88 | 0.98 |
| 2 | B*27:05 | 11 | -0.41 | 0.48 | 0.55 | 4 | -0.23 | 0.8 | -- |
| 3 | B*14 | 24 | -0.37 | 0.31 | 0.55 | 19 | -0.13 | 0.76 | 0.98 |
| 4 | C*08:02 | 23 | -0.3 | 0.41 | 0.55 | 17 | 0.01 | 0.98 | 0.98 |
| 5 | B*52 | 3 | -1.1 | 0.3 | -- | 2 | -1.7 | 0.19 | -- |
| 6 | A*25 | 7 | -0.81 | 0.25 | 0.55 | 5 | -0.99 | 0.23 | -- |
| 7 | B*35 | 46 | 0.37 | 0.2 | 0.55 | 32 | 0.35 | 0.28 | 0.56 |
| 8 | C*07 | 107 | -0.2 | 0.35 | 0.55 | 74 | -0.35 | 0.19 | 0.56 |
| 9 | Composite Variable | N/A | -0.26 | 0.064 | 0.51 | N/A | -0.19 | 0.25 | 0.56 |

^a^ Alleles 1-6 were previously designated as “protective” alleles, while alleles 7-8 were defined as “risk” alleles, in the International HIV Controllers genomewide association study (16). The “Composite Variable” combines each of the above alleles, counting each protective allele as +1 and each risk allele as -1.

^b^ We list minor allele count (MAC), rather than MAF, due to the multiallelic nature of these variants. Alleles with MAC<=5 are greyed out to highlight low observed frequencies in our study population.

^c^ Beta estimate from multivariate linear mixed model adjusted for age, sex, nadir CD4+ T cell count, timing of ART initiation, and ancestry.

^d^ Two-sided p-values; p<0.05 are annotated with asterisks.

^e^ Two-sided false discovery rate (FDR)-adjusted Benjamini-Hochberg q-values.

**Table S6.** HLA associations with HIV intact DNA (copies/10^6^ CD4+ T cells).

| **HIV Intact DNA** | | | | | | | | | |
| --- | --- | --- | --- | --- | --- | --- | --- | --- | --- |
|  |  | **Full Cohort** | | | | **European Subgroup** | | | |
|  | HLA Allele^a^ | MAC^b^ | Beta^c^ | p^d^ | q^e^ | MAC^b^ | Beta^c^ | p^d^ | q^e^ |
| 1 | B*57:01 | 9 | -0.59 | 0.24 | 0.77 | 8 | -0.42 | 0.43 | 0.76 |
| 2 | B*27:05 | 11 | -0.48 | 0.29 | 0.77 | 4 | -1.4 | 0.057 | -- |
| 3 | B*14 | 20 | 0.072 | 0.82 | 0.9 | 16 | -0.11 | 0.76 | 0.76 |
| 4 | C*08:02 | 19 | 0.039 | 0.9 | 0.9 | 15 | -0.15 | 0.67 | 0.76 |
| 5 | B*52 | 3 | -1.1 | 0.21 | -- | 2 | -1.2 | 0.25 | -- |
| 6 | A*25 | 6 | -1.2 | 0.054 | 0.43 | 4 | -1.5 | *0.044 | -- |
| 7 | B*35 | 43 | -0.069 | 0.78 | 0.9 | 29 | 0.21 | 0.46 | 0.76 |
| 8 | C*07 | 101 | -0.09 | 0.61 | 0.9 | 69 | 0.1 | 0.66 | 0.76 |
| 9 | Composite Variable | N/A | -0.1 | 0.4 | 0.8 | N/A | -0.25 | 0.079 | 0.47 |

^a^ Alleles 1-6 were previously designated as “protective” alleles, while alleles 7-8 were defined as “risk” alleles, in the International HIV Controllers genomewide association study (16). The “Composite Variable” combines each of the above alleles, counting each protective allele as +1 and each risk allele as -1.

^b^ We list minor allele count (MAC), rather than MAF, due to the multiallelic nature of these variants. Alleles with MAC<=5 are greyed out to highlight low observed frequencies in our study population.

^c^ Beta estimate from multivariate linear mixed model adjusted for age, sex, nadir CD4+ T cell count, timing of ART initiation, and ancestry.

^d^ Two-sided p-values; p<0.05 are annotated with asterisks.

^e^ Two-sided false discovery rate (FDR)-adjusted Benjamini-Hochberg q-values.

**Table S7.** Full list of single nucleotide polymorphisms (SNPs) from Table 1 that were associated with HIV total DNA (upper table) and unspliced RNA (lower table) in the total study population, ranked by p-value. Benjamini-Hochberg false discovery rate (FDR) adjusted q<0.05 shown in bold font.

| **SNP** | **Chrom** | **Position** | **Nearest Gene** | **Gene Location** | **MAF^a^** | **Beta^b^** | **eQTL Effect^c^** | **PVE^d^** | **p^e^** | **q^f^** | **Description** |
| --- | --- | --- | --- | --- | --- | --- | --- | --- | --- | --- | --- |
| **HIV TOTAL DNA** | | | | | | | | | | | |
| **Full Cohort** | | | | | | | | | | | |
| rs34814968 | chr21 | 41421873 | *MX1* | 5’UTR^g^ | 0.45 | -1.2 | -0.26 | 0.15 | 1.3x10^-7^ | **0.02** | Antiviral. Upregulated in HIV+ compared to uninfected individuals (63) and associated with higher viremia among HIV+ individuals (64). Associated with HIV-1 latency (61). MX2, but not MX1, was shown to be capable of directly acting against HIV-1 virus (58-59). |
| rs459482 | chr21 | 41421864 | *MX1* | 5’UTR^g^ | 0.45 | -1.2 | -0.26 | 0.15 | 1.3x10^-7^ | **0.02** |  |
| rs74867009 | chr12 | 25063777 | *LRMP* | 5’UTR^g^ | 0.06 | -2.5 |  | 0.15 | 1.5x10^-7^ | **0.02** | Delivers peptides to MHC class I molecules. (65) Differentially expressed in lymphatic tissue (66) and peripheral blood mononuclear cells (PBMCs) (67) in HIV+ individuals in response to ART initiation and cessation, respectively. |
| rs469073 | chr21 | 41444274 | *MX1* | Intronic | 0.50 | 1.1 |  | 0.13 | 5.8x10^-7^ | **0.02** |  |
| rs462687 | chr21 | 41425130 | *MX1* | Intronic | 0.51 | -1.1 | -0.26 | 0.13 | 8.4x10^-7^ | **0.02** |  |
| rs364105 | chr21 | 41446429 | *MX1* | Intronic | 0.51 | 1.1 |  | 0.13 | 8.7x10^-7^ | **0.02** |  |
| rs364203 | chr21 | 41446430 | *MX1* | Intronic | 0.51 | 1.1 |  | 0.13 | 8.7x10^-7^ | **0.02** |  |
| rs469483 | chr21 | 41446588 | *MX1* | Intronic | 0.51 | 1.1 |  | 0.13 | 8.7x10^-7^ | **0.02** |  |
| rs364199 | chr21 | 41446205 | *MX1* | Intronic | 0.51 | 1.1 |  | 0.13 | 9.5x10^-7^ | **0.02** |  |
| rs363882 | chr21 | 41446346 | *MX1* | Intronic | 0.51 | 1.1 |  | 0.13 | 9.5x10^-7^ | **0.02** |  |
| rs469390 | chr21 | 41446003 | *MX1* | Exonic | 0.51 | 1.1 |  | 0.13 | 9.5x10^-7^ | **0.02** |  |
| rs364143 | chr21 | 41447005 | *MX1* | Intronic | 0.51 | 1.1 |  | 0.13 | 9.5x10^-7^ | **0.02** |  |
| rs364043 | chr21 | 41447022 | *MX1* | Intronic | 0.51 | 1.1 |  | 0.13 | 9.5x10^-7^ | **0.02** |  |
| rs369454 | chr21 | 41447036 | *MX1* | Intronic | 0.51 | 1.1 |  | 0.13 | 9.5x10^-7^ | **0.02** |  |
| rs365628 | chr21 | 41447039 | *MX1* | Intronic | 0.51 | 1.1 |  | 0.13 | 9.5x10^-7^ | **0.02** |  |
| rs467773 | chr21 | 41446179 | *MX1* | Intronic | 0.51 | 1.1 |  | 0.13 | 9.5x10^-7^ | **0.02** |  |
| rs467960 | chr21 | 41440964 | *MX1* | Exonic | 0.47 | 1.1 | 0.13 | 0.12 | 1.2x10^-6^ | **0.02** |  |
| rs464138 | chr21 | 41426246 | *MX1* | 5’UTR^g^ | 0.41 | -1.0 | -0.27 | 0.12 | 1.5x10^-6^ | **0.02** |  |
| rs469383 | chr21 | 41445738 | *MX1* | Intronic | 0.51 | 1.1 |  | 0.12 | 1.5x10^-6^ | **0.02** |  |
| rs464783 | chr21 | 41424553 | *MX1* | Intronic | 0.40 | -1.1 | -0.28 | 0.12 | 1.5x10^-6^ | **0.02** |  |
| rs459498 | chr21 | 41423100 | *MX1* | Intronic | 0.40 | -1.1 | -0.28 | 0.12 | 1.5x10^-6^ | **0.02** |  |
| rs461981 | chr21 | 41422154 | *MX1* | Intronic | 0.40 | -1.1 | -0.28 | 0.12 | 1.5x10^-6^ | **0.02** |  |
| rs148214 | chr21 | 41442291 | *MX1* | Intronic | 0.48 | 1.1 | 0.12 | 0.12 | 1.7x10^-6^ | **0.02** |  |
| rs467987 | chr21 | 41442345 | *MX1* | Intronic | 0.48 | 1.1 | 0.12 | 0.12 | 1.7x10^-6^ | **0.02** |  |
| rs468704 | chr21 | 41442549 | *MX1* | Intronic | 0.48 | 1.1 | 0.12 | 0.12 | 1.7x10^-6^ | **0.02** |  |
| rs468440 | chr21 | 41442594 | *MX1* | Intronic | 0.48 | 1.1 | 0.12 | 0.12 | 1.7x10^-6^ | **0.02** |  |
| rs468747 | chr21 | 41442939 | *MX1* | Intronic | 0.48 | 1.1 | 0.12 | 0.12 | 1.7x10^-6^ | **0.02** |  |
| rs468414 | chr21 | 41443367 | *MX1* | Intronic | 0.48 | 1.1 | 0.12 | 0.12 | 1.7x10^-6^ | **0.02** |  |
| rs10670165 | chr21 | 41443170 | *MX1* | Intronic | 0.48 | 1.1 | 0.12 | 0.12 | 1.7x10^-6^ | **0.02** |  |
| rs364197 | chr21 | 41443442 | *MX1* | Intronic | 0.48 | 1.1 | 0.12 | 0.12 | 1.7x10^-6^ | **0.02** |  |
| rs468786 | chr21 | 41443293 | *MX1* | Intronic | 0.48 | 1.1 | 0.12 | 0.12 | 1.8x10^-6^ | **0.02** |  |
| rs1638369 | chr21 | 41444193 | *MX1* | Intronic | 0.48 | 1.1 | 0.12 | 0.12 | 1.8x10^-6^ | **0.02** |  |
| rs462698 | chr21 | 41444139 | *MX1* | Intronic | 0.48 | 1.1 |  | 0.12 | 1.8x10^-6^ | **0.02** |  |
| rs469012 | chr21 | 41443535 | *MX1* | Intronic | 0.48 | 1.1 | 0.12 | 0.12 | 1.8x10^-6^ | **0.02** |  |
| rs456018 | chr21 | 41421797 | *MX1* | Intronic | 0.40 | -1.1 | -0.28 | 0.12 | 2.1x10^-6^ | **0.02** |  |
| rs35477708 | chr21 | 41446620 | *MX1* | Intronic | 0.48 | 1.0 |  | 0.12 | 2.8x10^-6^ | **0.02** |  |
| rs469270 | chr21 | 41445250 | *MX1* | Intronic | 0.48 | 1.0 | 0.12 | 0.12 | 2.8x10^-6^ | **0.02** |  |
| rs467537 | chr21 | 41447082 | *MX1* | Intronic | 0.48 | 1.1 |  | 0.11 | 3.1x10^-6^ | **0.03** |  |
| rs469093 | chr21 | 41444578 | *MX1* | Intronic | 0.48 | 1.1 | 0.12 | 0.11 | 3.1x10^-6^ | **0.03** |  |
| rs457274 | chr21 | 41420558 | *MX1* | 5’UTR^g^ | 0.41 | -1.0 | -0.27 | 0.11 | 3.2x10^-6^ | **0.03** |  |
| rs467574 | chr21 | 41447278 | *MX1* | Intronic | 0.41 | 1.1 | 0.12 | 0.11 | 3.7x10^-6^ | **0.03** |  |
| rs751660317 | chr2 | 28786774 | *PPP1CB* | Intronic | 0.07 | -1.8 |  | 0.11 | 4.1x10^-6^ | **0.03** | Encodes a subunit of PP1. Depletion of PPP1CB was shown to reduce the antiviral potency of MX2 against HIV (60). PP1 is involved in transcription of HIV-1; inhibition of PP1 inhibits HIV-1 transcription (68). |
| rs469288 | chr21 | 41445655 | *MX1* |  | 0.52 | 1.0 |  | 0.11 | 7.2x10^-6^ | **0.05** |  |
| rs776025235 | chr6 | 51638799 | *PKHD1* | Intronic | 0.09 | -1.7 |  | 0.09 | 2.5x10^-5^ | 0.17 | Predicted to have a transmembrane-spanning domain and an immunoglobulin-like plexin-transcription-factor domain. We could not find a direct relationship with HIV in the literature. |
| N/A | chrX | 41382082 | *DDX3X; NYX* | Intergenic | 0.18 | -1.4 |  | 0.09 | 2.5x10^-5^ | 0.17 | *DDX3X* regulates the production of type I IFNs (69) and encodes a protein that shuttles HIV-1 RNA from the nucleus to the cytoplasm. Is upregulated in HIV-infected cells; knockdown of *DDX3X* suppresses HIV-1 replication (70). Also plays a key role in innate antimicrobial immunity (69). |
| rs468259 | chr21 | 41453222 | *MX1* | Intronic | 0.48 | 0.9 |  | 0.09 | 2.6x10^-5^ | 0.18 |  |
| rs17506750 | chr14 | 32599402 | *AKAP6* | Intronic | 0.07 | -1.8 |  | 0.09 | 3.0x10^-5^ | 0.20 | Binds to regulatory subunits of protein kinase A (PKA) and anchors them to the nuclear membrane. PKA activation has been associated with HIV-1 infection, T cell proliferation, and dysfunction (71-72). |
| rs469066 | chr21 | 41444089 | *MX1* | Intronic | 0.64 | 1.0 | 0.24 | 0.09 | 3.3x10^-5^ | 0.22 |  |
| rs463856 | chr21 | 41431931 | *MX1* | Intronic | 0.64 | -1.0 |  | 0.09 | 3.6x10^-5^ | 0.23 |  |
| **European Ancestry Subgroup** | | | | | | | | | | | |
| rs469483 | chr21 | 41446588 | *MX1* | Intron/Exon/  5’UTR^g^ | 0.55 | 1.3 |  | 0.20 | 9.2x10^-7^ | **0.02** | Antiviral. Upregulated in HIV+ compared to uninfected individuals (63) and associated with higher viremia among HIV+ individuals (64). Associated with HIV-1 latency (61). MX2, but not MX1, was shown to be capable of directly acting against HIV-1 virus (58-59). |
| rs364203 | chr21 | 41446430 | *MX1* | Intronic | 0.55 | 1.3 |  | 0.20 | 9.2x10^-7^ | **0.02** |  |
| rs469288 | chr21 | 41445655 | *MX1* | Intronic | 0.55 | 1.3 |  | 0.20 | 9.2x10^-7^ | **0.02** |  |
| rs364105 | chr21 | 41446429 | *MX1* | Intronic | 0.55 | 1.3 |  | 0.20 | 9.2x10^-7^ | **0.02** |  |
| rs369454 | chr21 | 41447036 | *MX1* | Intronic | 0.54 | 1.3 |  | 0.20 | 1.0x10^-6^ | **0.02** |  |
| rs364043 | chr21 | 41447022 | *MX1* | Intronic | 0.54 | 1.3 |  | 0.20 | 1.0x10^-6^ | **0.02** |  |
| rs364143 | chr21 | 41447005 | *MX1* | Intronic | 0.54 | 1.3 |  | 0.20 | 1.0x10^-6^ | **0.02** |  |
| rs469073 | chr21 | 41444274 | *MX1* | Intronic | 0.54 | 1.3 |  | 0.20 | 1.0x10^-6^ | **0.02** |  |
| rs365628 | chr21 | 41447039 | *MX1* | Intronic | 0.54 | 1.3 |  | 0.20 | 1.0x10^-6^ | **0.02** |  |
| rs363882 | chr21 | 41446346 | *MX1* | Intronic | 0.54 | 1.3 |  | 0.20 | 1.0x10^-6^ | **0.02** |  |
| rs364199 | chr21 | 41446205 | *MX1* | Intronic | 0.54 | 1.3 |  | 0.20 | 1.0x10^-6^ | **0.02** |  |
| rs467773 | chr21 | 41446179 | *MX1* | Intronic | 0.54 | 1.3 |  | 0.20 | 1.0x10^-6^ | **0.02** |  |
| rs469390 | chr21 | 41446003 | *MX1* | Exonic | 0.54 | 1.3 |  | 0.20 | 1.0x10^-6^ | **0.02** |  |
| rs469383 | chr21 | 41445738 | *MX1* | Intronic | 0.54 | 1.2 |  | 0.19 | 1.7x10^-6^ | **0.04** |  |
| rs34814968 | chr21 | 41421873 | *MX1* | 5’UTR^g^ | 0.46 | -1.2 | -0.26 | 0.18 | 5.0x10^-6^ | 0.08 |  |
| rs459482 | chr21 | 41421864 | *MX1* | 5’UTR^g^ | 0.46 | -1.2 | -0.26 | 0.18 | 5.0x10^-6^ | 0.08 |  |
| . | chr21 | 41424232 | *MX1* | Intronic | 0.39 | -1.2 |  | 0.18 | 5.4x10^-6^ | 0.08 |  |
| rs464138 | chr21 | 41426246 | *MX1* | 5’UTR^g^ | 0.44 | -1.1 | -0.27 | 0.17 | 5.9x10^-6^ | 0.08 |  |
| rs457274 | chr21 | 41420558 | *MX1* | 5’UTR^g^ | 0.44 | -1.1 | -0.27 | 0.17 | 7.8x10^-6^ | 0.08 |  |
| rs468414 | chr21 | 41443367 | *MX1* | Intronic | 0.53 | 1.1 | 0.12 | 0.17 | 7.9x10^-6^ | 0.08 |  |
| rs35477708 | chr21 | 41446620 | *MX1* | Intronic | 0.53 | 1.1 |  | 0.17 | 7.9x10^-6^ | 0.08 |  |
| rs364197 | chr21 | 41443442 | *MX1* | Intronic | 0.53 | 1.1 | 0.12 | 0.17 | 7.9x10^-6^ | 0.08 |  |
| rs469270 | chr21 | 41445250 | *MX1* | Intronic | 0.53 | 1.1 | 0.12 | 0.17 | 7.9x10^-6^ | 0.08 |  |
| rs148214 | chr21 | 41442291 | *MX1* | Intronic | 0.53 | 1.1 | 0.12 | 0.17 | 7.9x10^-6^ | 0.08 |  |
| rs10670165 | chr21 | 41443170 | *MX1* | Intronic | 0.53 | 1.1 | 0.12 | 0.17 | 7.9x10^-6^ | 0.08 |  |
| rs468747 | chr21 | 41442939 | *MX1* | Intronic | 0.53 | 1.1 | 0.12 | 0.17 | 7.9x10^-6^ | 0.08 |  |
| rs468440 | chr21 | 41442594 | *MX1* | Intronic | 0.53 | 1.1 | 0.12 | 0.17 | 7.9x10^-6^ | 0.08 |  |
| rs467987 | chr21 | 41442345 | *MX1* | Intronic | 0.53 | 1.1 | 0.12 | 0.17 | 7.9x10^-6^ | 0.08 |  |
| rs468704 | chr21 | 41442549 | *MX1* | Intronic | 0.53 | 1.1 | 0.12 | 0.17 | 7.9x10^-6^ | 0.08 |  |
| rs468786 | chr21 | 41443293 | *MX1* | Intronic | 0.52 | 1.1 | 0.12 | 0.17 | 9.0x10^-6^ | 0.08 |  |
| rs467960 | chr21 | 41440964 | *MX1* | Exonic | 0.52 | 1.1 | 0.13 | 0.17 | 9.0x10^-6^ | 0.08 |  |
| rs469012 | chr21 | 41443535 | *MX1* | Intronic | 0.52 | 1.1 | 0.12 | 0.17 | 9.0x10^-6^ | 0.08 |  |
| rs462698 | chr21 | 41444139 | *MX1* | Intronic | 0.52 | 1.1 |  | 0.17 | 9.0x10^-6^ | 0.08 |  |
| rs467537 | chr21 | 41447082 | *MX1* | Intronic | 0.52 | 1.1 |  | 0.17 | 9.0x10^-6^ | 0.08 |  |
| rs1638369 | chr21 | 41444193 | *MX1* | Intronic | 0.52 | 1.1 | 0.12 | 0.17 | 9.0x10^-6^ | 0.08 |  |
| rs469093 | chr21 | 41444578 | *MX1* | Intronic | 0.52 | 1.1 | 0.12 | 0.17 | 9.0x10^-6^ | 0.08 |  |
| rs462687 | chr21 | 41425130 | *MX1* | Intronic | 0.46 | -1.1 | -0.26 | 0.16 | 1.1x10^-5^ | 0.09 |  |
| rs467574 | chr21 | 41447278 | *MX1* | Intronic | 0.50 | 1.1 | 0.12 | 0.16 | 1.3x10^-5^ | 0.11 |  |
| rs456018 | chr21 | 41421797 | *MX1* | Intronic | 0.44 | -1.1 | -0.28 | 0.16 | 1.8x10^-5^ | 0.14 |  |
| rs461981 | chr21 | 41422154 | *MX1* | Intronic | 0.44 | -1.1 | -0.28 | 0.16 | 1.9x10^-5^ | 0.14 |  |
| rs464783 | chr21 | 41424553 | *MX1* | Intronic | 0.44 | -1.1 | -0.28 | 0.16 | 1.9x10^-5^ | 0.14 |  |
| rs459498 | chr21 | 41423100 | *MX1* | Intronic | 0.44 | -1.1 | -0.28 | 0.16 | 1.9x10^-5^ | 0.14 |  |
| N/A | chr11 | 59574427 | *OSBP* | 3’UTR^g^ | 0.15 | -1.8 |  | 0.15 | 1.9x10^-5^ | 0.14 | Oxysterol binding protein involved in intracellular lipid transport. Associated with *in vitro* HIV-1 infection of monocyte-derived macrophages (MDMs) from highly (HIV)-exposed seronegatives (HESNs) (73). Is required for the replication of several human viruses such as hepatitis C (HCV), encephalomyocarditis (EMCV), Zika, etc. (74). |
| **HIV UNSPLICED RNA** | | | | | | | | | | | |
| **Full Cohort** | | | | | | | | | | | |
| N/A | chr19 | 17374631 | *PLVAP* | Intronic | 0.10 | -1.2 |  | 0.14 | 6.9x10^-7^ | 0.21 | Endothelial membrane protein, also controls the entry of lymphocytes and antigens into lymph nodes (75). Located within 30kb of *BST2* (aka tetherin), which is known to inhibit HIV-1 release by directly tethering virions to cells (76). |

^a^ MAF = minor allele frequency in the study population.

^b^ Beta = estimate from multivariate linear mixed model adjusted for age, sex, nadir CD4+ T cell count, timing of ART initiation, and ancestry.

^c^ eQTL effect = The normalized effect of the SNP on gene expression from previously reported human whole blood experimental data (57).

^d^ PVE = proportion of phenotype variance explained.

^e^ p = two sided p-value.

^f^ q = two-sided false discovery rate (FDR) Benjamini-Hochberg q-value.

^g^ UTR= untranslated region

**Table S8**. Host gene sets associated with HIV reservoir size using the Gene Ontology (GO) Biological Processes database from the multi-SNP rare variant gene-based analyses. Core enrichment genes contributing to q-values are shown in bold font.

| **Gene Set** | **NES^a^** | **q^b^** | **List of Genes in Gene Set** |
| --- | --- | --- | --- |
| **HIV TOTAL DNA** | | | |
| **Full Cohort** | | | |
| N Glycan Processing | 1.76 | 0.12 | ***MGAT4B, MAN2A1, PRKCSH, GNPTAB, ST8SIA3, MAN1A1****, MAN1C1, MGAT4A, FUT8, GANAB, ST8SIA2, MAN1B1, MAN1A2, ST8SIA4, MAN2A2* |
| **HIV UNSPLICED RNA** | | | |
| **Full Cohort** | | | |
| Single Stranded Viral RNA Replication VIA Double Stranded DNA Intermediate | 1.73 | 0.16 | ***APOBEC3D, APOBEC3C, TRIM28, APOBEC3F, TOP2B, APOBEC3H, APOBEC3G****, HMGA2, TASOR, RESF1, INPP5K, CXCL8, TOP2A, MPHOSPH8, AICDA, SETDB1, MORC2* |
| **HIV INTACT DNA** | | | |
| **Full Cohort** | | | |
| Linoleic Acid Metabolic Process | 1.71 | 0.22 | ***ALOX15B, ELOVL1, ALOX12B, ALOXE3, CYP2J2, GSTA1, ACSL1****, ELOVL5, GSTM2, ALOX12, GSTP1, ABCD1, FADS2, ELOVL3, FADS1, PNPLA8, ELOVL2* |
| Replacement Ossification | 1.69 | 0.18 | ***SMPD3, RUNX2, DLX5, MEF2C, BMP4, INPPL1, PHOSPHO1, PEX7, MMP16, FGF18, COL1A1****, TEK, COL2A1, GNAS, FGFR3, NAB2, BMP6, ALPL, COL13A1, FOXC1, IMPAD1, SCX, MEF2D, MMP14, MMP13, CSGALNACT1, NAB1* |
| Regulation of Peptidyl Serine Phosphorylation of STAT Protein | 1.67 | 0.19 | ***IFNA4, IFNA16, IFNA10, IFNA7, IFNA17, IFNA21, RET, IFNA14, IFNW1, LIF****, IFNB1, ERCC6, IFNA5, IFNK, IFNA8, IFNG, IFNA13, IFNA6, IFNE, IFNA2, PWP1, IFNA1* |
| Response to Lead Ion | 1.65 | 0.21 | ***PPP2CA, ATP7A, ANXA5, PPP5C****, FECH, CDK4, PPP1CA, SLC6A1, BECN1, PPP2CB, PTH, SLC34A1, QDPR, ALAD, BACE1, STAR, SPARC, APP, MAP1LC3A, PTGS2, CLDN1, DNMT3A, CAT* |
| Serine Phosphorylation of STAT Protein | 1.63 | 0.25 | ***IFNA4, IFNA16, IFNA10, IFNA7, IFNA17, IFNA21, RET, IFNA14, IFNW1, CDK5R1, LIF****, IFNB1, IL24, ERCC6, IFNA5, IFNK, CDK5, IFNA8, IFNG, IFNA13, IFNA6, IFNE, IFNA2, PWP1, IFNA1, NLK* |
| **European Ancestry Subgroup** | | | |
| Negative Regulation of Viral Entry into Host Cell | 1.81 | **0.03** | ***IFITM1, IFITM3, APCS, IFITM2, FCN3, FCN1, GSN, TRIM59****, SNX3, TRIM25, PTX3, TRIM11, TRIM8, MID2, TRIM5, IFNA2* |

^a^ NES = Normalized enrichment score = gene set enrichment score normalized based on the number of genes in the gene set.

^b^ q = two-sided false discovery rate (FDR) Benjamini-Hochberg q<0.05 shown in bold font.

**Table S9.** As an example of the lack of overlap in SNP associations by ancestral subgroups: the top ten SNPs associated with HIV total DNA are shown for the non-European (upper table) and the African-American (bottom table) ancestry subgroups. These results compare to the SNPs associated with HIV total DNA in the total population and European ancestry subgroup, as shown in **Table 1**.

| **SNP** | **Chrom** | **Position** | **Nearest Gene** | **Gene Location** | **MAF^a^** | **Beta^b^** | **PVE^c^** | **p^d^** | **q^e^** | **Rank^f^ Total** | **Rank^f^ EUR** |
| --- | --- | --- | --- | --- | --- | --- | --- | --- | --- | --- | --- |
| **HIV TOTAL DNA** | | | | | | | | | | | |
| **Non-European Ancestry Subgroup** | | | | | | | | | | | |
| NA^g^ | chr17 | 29439869 | *TAOK1* | Intron | 0.08 | -3.4 | 0.34 | 8.0x10^-6^ | 0.99 | 1393 | 180210 |
| rs10770125 | chr11 | 2147784 | *INS-IGF2* | Exon | 0.43 | -1.9 | 0.30 | 2.2x10^-5^ | 0.99 | 12114 | 149522 |
| rs3743598 | chr16 | 75612787 | *ADAT1* | Exon | 0.59 | 1.8 | 0.28 | 4.1x10^-5^ | 0.99 | 10296 | 168996 |
| NA^g^ | chr19 | 15194957 | *NOTCH3* | Intron | 0.38 | -2.5 | 0.28 | 4.3x10^-5^ | 0.99 | 1806 | 79122 |
| rs1003483 | chr11 | 2146313 | *IGF2-AS* | Intron/Exon | 0.42 | -1.8 | 0.27 | 5.8x10^-5^ | 0.99 | 16656 | 149522 |
| NA^g^ | chr17 | 29439872 | *TAOK1* | Intron | 0.07 | -3.4 | 0.27 | 7.2x10^-5^ | 0.99 | 5082 | 69970 |
| NA^g^ | chr17 | 29439873 | *TAOK1* | Intron | 0.07 | -3.3 | 0.27 | 7.5x10^-5^ | 0.99 | 5130 | 73764 |
| NA^g^ | chr17 | 29439871 | *TAOK1* | Intron | 0.07 | -3.3 | 0.27 | 7.5x10^-5^ | 0.99 | 4723 | 61379 |
| rs17659204 | chr2 | 6931128 | *RNF144A* | Intron | 0.14 | -2.4 | 0.26 | 8.7x10^-5^ | 0.99 | 5351 | 300439 |
| rs751660317 | chr2 | 28786774 | *PPP1CB* | Intron | 0.10 | -2.2 | 0.25 | 1.1x10^-4^ | 0.99 | 42 | 565 |
| **African-American Ancestry Subgroup** | | | | | | | | | | | |
| rs6946155 | chr7 | 5299139 | *SLC29A4* | Intron | 0.28 | -3.3 | 0.75 | 4.1x10^-4^ | 0.98 | 126566 | 28785 |
| rs770098394 | chr15 | 43635425 | *PPIP5K1P1-CATSPER2* | Intron | 0.10 | -4.0 | 0.75 | 4.4x10^-4^ | 0.98 | 43050 | NA |
| rs372669041 | chr17 | 45242067 | *FMNL1* | Exon | 0.54 | -2.3 | 0.73 | 5.0x10^-4^ | 0.98 | NA | NA |
| rs7250831 | chr19 | 55388066 | *RPL28* | Intron/Exon | 0.10 | -3.8 | 0.72 | 5.6x10^-4^ | 0.98 | NA | NA |
| rs780679191 | chrX | 156025084 | *WASH6P* | Exon | 0.16 | -3.3 | 0.70 | 6.3x10^-4^ | 0.98 | NA | NA |
| rs34551990 | chr7 | 5300371 | *SLC29A4* | Intron | 0.26 | -3.4 | 0.70 | 6.9x10^-4^ | 0.98 | 67704 | 94946 |
| rs62378453 | chr5 | 141408283 | *PCDHGB6* | Intron/Exon | 0.20 | -3.3 | 0.69 | 7.0x10^-4^ | 0.98 | 14391 | 127130 |
| rs7224464 | chr17 | 4449244 | *SPNS3* | Exon | 0.48 | 2.4 | 0.69 | 7.2x10^-4^ | 0.98 | 177961 | 147106 |
| rs1752435 | chr14 | 59479376 | *L3HYPDH* | Intron | 0.14 | -3.7 | 0.69 | 7.2x10^-4^ | 0.98 | 31937 | 215209 |
| rs373114591 | chr9 | 39102618 | *CNTNAP3* | Exon | 0.10 | -3.8 | 0.69 | 7.2x10^-4^ | 0.98 | NA | NA |

^a^ MAF = minor allele frequency in the study population.

^b^ Beta = estimate from multivariate linear mixed model adjusted for age, sex, nadir CD4+ T cell count, timing of ART initiation, and ancestry.

^c^ PVE = proportion of phenotype variance explained.

^d^ p = two sided p-value.

^e^ q = two-sided false discovery rate (FDR) Benjamini-Hochberg q-value.

^f^ Rank = SNP “rank” (from lowest to highest two-sided q-value) for the total population (“Total”) or for the European ancestry subgroup (“EUR”). Bold font represents statistically significant SNP(s) for that subgroup at a two-sided q<0.05. “NA” stated if the MAF is below 5% in the reference population.

^g^ NA = novel variant; no rs number.
